# Systematic elucidation of genetic mechanisms underlying cholesterol uptake

**DOI:** 10.1101/2023.01.09.500804

**Authors:** Marisa C. Hamilton, James D. Fife, Ersin Akinci, Tian Yu, Benyapa Khowpinitchai, Minsun Cha, Sammy Barkal, Thi Tun Thi, Grace H.T. Yeo, Juan Pablo Ramos Barroso, Matthew Jake Francoeur, Minja Velimirovic, David K. Gifford, Guillaume Lettre, Haojie Yu, Christopher A. Cassa, Richard I. Sherwood

## Abstract

Genetic variation contributes greatly to LDL cholesterol (LDL-C) levels and coronary artery disease risk. By combining analysis of rare coding variants from the UK Biobank and genome-scale CRISPR-Cas9 knockout and activation screening, we have substantially improved the identification of genes whose disruption alters serum LDL-C levels. We identify 21 genes in which rare coding variants significantly alter LDL-C levels at least partially through altered LDL-C uptake. We use co-essentiality-based gene module analysis to show that dysfunction of the RAB10 vesicle transport pathway leads to hypercholesterolemia in humans and mice by impairing surface LDL receptor levels. Further, we demonstrate that loss of function of *OTX2* leads to robust reduction in serum LDL-C levels in mice and humans by increasing cellular LDL-C uptake. Altogether, we present an integrated approach that improves our understanding of genetic regulators of LDL-C levels and provides a roadmap for further efforts to dissect complex human disease genetics.

## Introduction

Coronary artery disease (CAD) is one of the most common causes of death and disability worldwide^1^. Genetic differences in cholesterol metabolism contribute substantially to CAD risk, along with well-known environmental and clinical risk factors. Serum cholesterol measurements are quantitative and nearly uniformly measured across biobanks, providing rich human phenotypic data that can aid in the identification of genetic risk factors for CAD. Rare monogenic coding variants have been found to dramatically alter low-density lipoprotein cholesterol (LDL-C) uptake, most notably in *LDLR, APOB*, and *PCSK9*^2^. Common, lower effect size genetic variants have also been identified through genome-wide association screens (GWAS) which significantly impact LDL-C levels and CAD risk^3–7^.

Unraveling the genetics of cholesterol is central to the development of therapeutics for heart disease. Genetic findings from rare and common variants with effects on cholesterol levels have led to the development of FDA-approved drugs targeting PCSK9 and drugs in the clinical pipeline targeting ANGPTL3, APOC3, and LPA^8,9^, among others. Additionally, statin drugs and ezetimibe, which are commonly prescribed to patients with elevated LDL-C levels and high CAD risk, inhibit HMGCR and NPC1L1 protein function respectively^10^, and human genetics studies have identified that variants within and surrounding the *HMGCR* and *NPC1L1* genes impact LDL-C levels^3,11–14^.

Nonetheless, the genetics of cholesterol levels are far from completely understood. A recent trans-ancestry GWAS meta-analysis from the Global Lipids Genetics Consortium (GLGC), using data from 1.65 million subjects, has identified >900 genome-wide significant loci associated with blood lipid levels, including >400 loci associated with LDL-C^15^. Yet, the vast majority of genes or non-coding RNAs which modulate these loci have not been identified. Direct analysis of associations between deleterious coding variants and cholesterol levels from large exome sequencing cohorts, including rare-variant burden analyses of up to 454,787 individuals in the UK Biobank (UKB) cohort, have found at most 14 genes for which coding variation is significantly associated with LDL-C levels^11–14^. Thus, a major disconnect currently remains between the hundreds of loci flagged in GWAS and the paucity of genes shown to drive these associations.

Organismal regulation of LDL-C is accomplished through a complex set of mechanisms. Cholesterol is absorbed from ingested food in the intestine and also produced through the cholesterol biosynthetic pathway primarily in the liver. Cholesterol is secreted into the bloodstream in chylomicrons and very-low-density lipoprotein (VLDL) and converted to LDL by lipoprotein lipase. Unmodified LDL is removed from the bloodstream by the LDL receptor (LDLR), primarily in the liver, while oxidized LDL can be absorbed by macrophage scavenger receptors or deposited in the arterial intima to form plaques. In spite of the complex nature of lipoprotein metabolism, liver cell LDL uptake has been shown to be a dominant mechanism controlling serum LDL-C levels, as it is the primary deficiency in familial hypercholesterolemia patients^16^ and the primary target of statins and PCSK9 inhibitors^17^.

Liver cell LDL uptake and cholesterol metabolism are amenable to dissection in a dish, and cell culture models have been instrumental in dissecting the sterol-responsive SREBP and LXR pathways^18,19^, the two major transcription regulators of cholesterol biosynthesis and LDL-C uptake machinery. Cellular uptake of fluorescent LDL-C has been used in the context of RNAi and CRISPR-Cas9 screening^20–23^, identifying genetic regulators of cholesterol metabolism. In particular, a recent genome-wide CRISPR-Cas9 screen in hepatic cell lines identified 163 genes whose mutation significantly alters LDL-C uptake^23^. Yet, how these *in vitro* LDL-C uptake regulators are functionally related in biological pathways, and whether variation in these genes plays a role in controlling serum LDL-C levels is poorly understood.

In this work, we combine human biobank coding variant burden analysis, CRISPR screening in human cell models, and gene network analysis to dissect the genetics of LDL-C uptake. We perform genome-scale CRISPR-Cas9 knockout screening for LDL-C uptake-altering genes and identify 490 genes whose mutation impacts liver cell LDL-C uptake. Through CRISPR gene activation screening, we identify genes whose overexpression and knockout alter LDL-C uptake in opposing directions and thus represent tunable regulators of LDL-C uptake. Through a co-essential gene network analysis pipeline, we further identify 33 gene modules significantly associated with LDL-C uptake.

To determine the disease relevance of the genes and pathways identified through our LDL-C uptake CRISPR screens, we examine LDL-C levels in carriers of rare coding variants within the UKB dataset, including through a coding burden approach that incorporates estimated deleteriousness of each variant. Using this approach, we find 21 genes that putatively alter LDL-C levels at least partially through altered LDL-C uptake, finding that our cellular LDL-C uptake CRISPR-Cas9 screens are significantly enriched with genes that have a coding burden. We also employ a co-essentiality module burden analysis, finding that genes associated with the GTPase RAB10 and exocytosis machinery greatly impact LDL-C levels, defining a novel hypercholesterolemia pathway. We demonstrate that RAB10 exocytosis pathway mutations impair trafficking of LDLR to the cell surface in human liver cells and that their loss of function leads to hypercholesterolemia in mice. Additionally, our integrated pipeline identifies several novel genes that significantly decrease serum LDL-C levels upon disruption at least in part through increased LDL-C uptake. We show that suppression of one such gene, *OTX2*, in mouse liver leads to robust decrease in serum LDL-C. In sum, our integrated approach evaluating the results of deleterious coding variants in human cell models through CRISPR screening and in the human population through exome burden analysis reveals novel genetic contributors to LDL-C levels and provides a roadmap to improved understanding of quantitative human trait genetics.

## Results

### Genome-scale CRISPR-Cas9-nuclease screening identifies hepatic cell LDL-C uptake-altering genes

Uptake of fluorescently labeled LDL-C by liver cells is a well-characterized assay for determining genetic contributors to LDL-C levels^20–24^. We have used a CRISPR-Cas9 screening platform to determine which human genes govern uptake of LDL-C in HepG2 hepatocellular carcinoma cells^25^, a well-studied model of liver cholesterol metabolism which has a relatively normal karyotype^26^ and has been the subject of among the most extensive epigenetic profiling through ENCODE^27^.

To assess the dynamic range of fluorescent LDL-C uptake after mutation of known LDL-C uptake-altering genes, we lentivirally infected HepG2 with gRNAs targeting *LDLR* and the LDLR-degrading E3 ubiquitin-protein ligase *MYLIP* (*a.k.a*. IDOL)^28^. We assessed LDL-C uptake in serum-starved (SS) and serum-containing (SER) conditions, find that Cas9-nuclease mutation of *LDLR* leads to a 3-4-fold reduction in LDL-C uptake (p<0.0001 in SS and SER) while mutation of *MYLIP* leads to up to 1.5-1.7-fold increase in LDL-C uptake (p<0.0001 in SS and SER, **Figure 1A, Figure S1A**).

**Figure 1:**
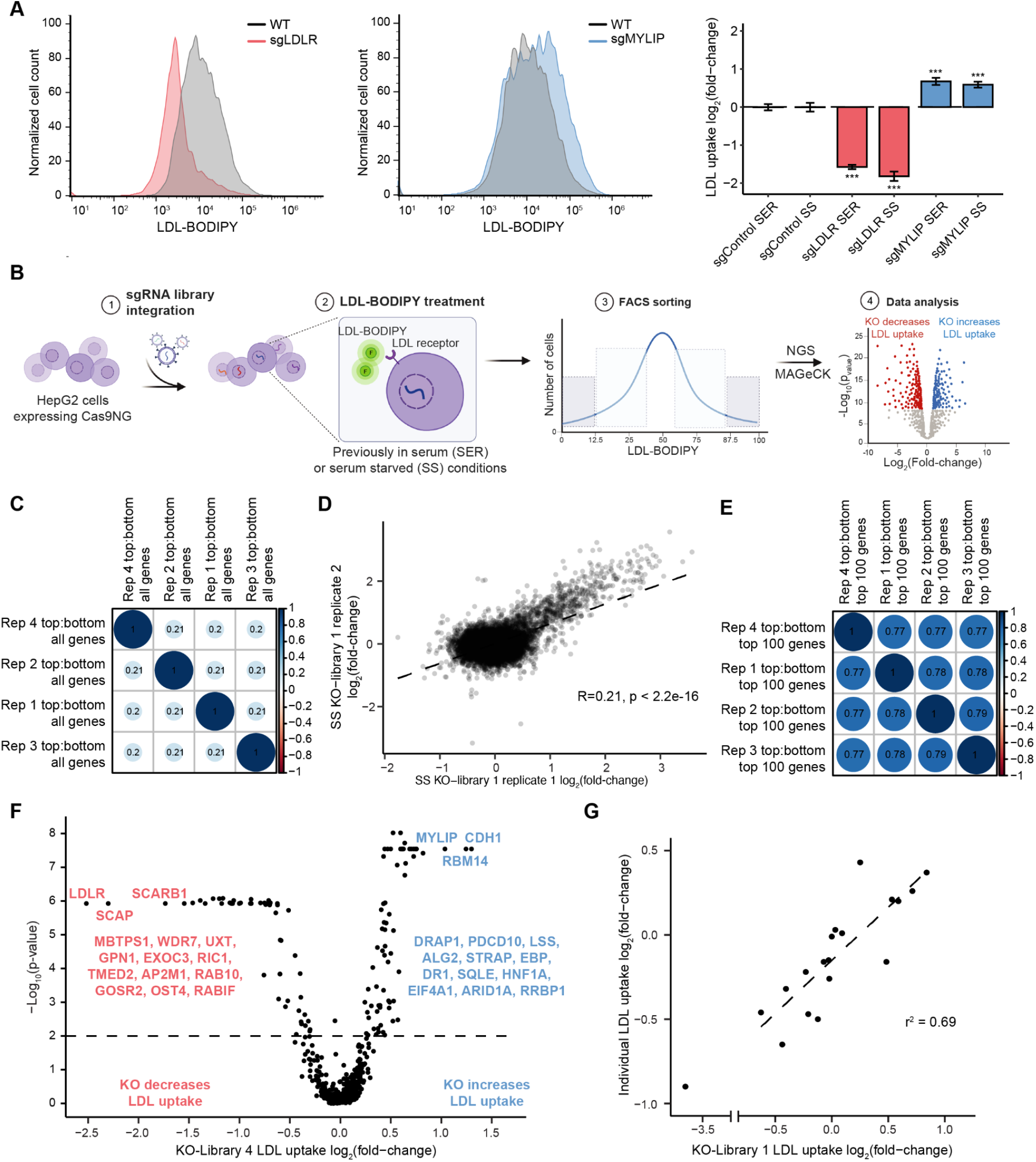
CRISPR screening identifies genes required for LDL-C uptake. A: Flow cytometric analysis of LDL-C uptake in HepG2 cells with CRISPR-Cas9 knockout of LDLR (red) and MYLIP (blue) in SER and SS conditions. n = 10. Two-tailed t-test. P-values are represented by asterisks (*p<0.05, **p≤0.01, ***p≤0.001). B: Schematic of LDL-C uptake CRISPR-Cas9 knockout screening. HepG2 constitutively expressing Cas9NG are incubated with a lentiviral gRNA library, followed by selection for infected cells. Cells are incubated in serum-containing or serum-starved conditions overnight followed by incubation with fluorescent LDL (BODIPY-LDL) for 4-6 hours followed by flow cytometric sorting of cells into 2-4 populations with most and least LDL-C uptake. Genomic DNA from sorted populations is prepared for nextgen sequencing (NGS), and gRNA counts are analyzed by MAGECK to identify genes associated with altered LDL-C uptake. C-E: Spearman correlation analysis of the ratio of representation of all gRNAs (C,D) or gRNAs in the 100 genes with strongest effects on LDL-C uptake (E) in cells with top 37.5% vs. bottom 37.5% LDL-C uptake in KO-Library 1 in SS conditions. F: Volcano plot showing the MAGECK LDL-C uptake log2-fold-change (x-axis) and minimum MAGECK −Log10-p-value (y-axis) for the 522 genes in KO-Library 4 in SS condition. Several significant genes are listed. Permutations = 100,000. G: Comparison of LDL-C uptake log2-fold-change for 20 genes between KO-Library 1 and individual gRNA knockout testing (y-axis).

To identify genes associated with LDL-C uptake, we performed a high-throughput CRISPR-Cas9-nuclease screen targeting genes adjacent to variants associated with serum LDL-C from a large multi-ancestry GWAS from the Million Veteran Program (MVP) cohort^3^. We identified 2,634 genes within 1 Mb of the 293 lead genome-wide significant MVP serum LDL-C variants. We designed a gRNA library (KO-Library 1) targeting each of these genes with four gRNAs predicted by the Cas9 indel prediction algorithm inDelphi^29^ to induce frameshifts in >80% of edited alleles as well as 200 non-targeting control gRNAs.

We performed lentiviral CRISPR-Cas9-nuclease screens using KO-Library 1 in four biological replicates. RNA-seq analysis of HepG2 in SER vs. SS conditions showed robust upregulation of a gene set associated with lipid biosynthesis (**Supplemental Table 1**), and so we performed screens in each condition to identify genes responsive to this known environmental variable, ensuring >500 cells per gRNA at all stages^30^. We flow cytometrically isolate four populations per replicate with the lowest 12.5%, next lowest 25%, highest 12.5%, and next highest 25% LDL uptake (**Figure 1B**), using NGS on all sorted populations followed by the MAGeCK RRA pipeline^31^ to identify genes with significant effects on LDL-C uptake.

We find strong replicability in pre-sort gRNA counts (R=0.79-0.91, **Figure S1B**), appreciable replicability in the ratio of gRNA representation in cells with the highest vs. lowest 37.5% LDL uptake (R=0.2-0.21, **Figure 1C,D**) and strong replicability among the 100 genes with the strongest effect on LDL-C uptake (R=0.77-0.79, **Figure 1E**). MAGECK analysis identifies 37 genes whose disruption significantly alters LDL-C uptake (FDR <0.1) in either SS or SER conditions (**Figure S1C, Supplemental Table 2**).

To ask whether the resolution to identify significant LDL-C uptake-altering genes is improved by increasing the number of cells screened per gRNA, we designed a smaller gRNA library targeting the 360 genes with most robust effect on LDL-C uptake in our original screen according to MAGECK (KO-Library 2). Screening KO-Library 2 using the same protocol as for KO-Library 1 identified 152 genes whose disruption significantly alters LDL-C uptake (FDR <0.1, **Figure S1D, Supplemental Table 2**) in either SS or SER conditions and yielded results that are highly concordant with the larger screen.

Given the added power to identify LDL-C uptake-altering genes using smaller gRNA libraries, we performed CRISPR-Cas9 screening using two additional gRNA libraries. One library contains four gRNAs targeting each of the 1,803 annotated human transcription factors^32,33^ (KO-Library 3) to increase the resolution to identify transcriptional regulators of LDL-C uptake (**Figure S1E, Supplemental Table 2**). The final library (KO-Library 4) contains four gRNAs targeting 522 genes with significant or near-significant effects on LDL-C uptake from the three previous screens as well as from a recently published genome-wide LDL-C uptake CRISPR-Cas9 screen performed in Huh7 cells^23^. Altogether, the four screens identify a total of 490 genes whose knockout significantly alters LDL-C uptake in either SER or SS conditions (299 decrease uptake, 180 increase uptake, 11 do both at FDR <0.1, **Figure 1F, Supplemental Table 2**), substantially increasing the number of such genes as compared to previous published screens^22,23,34,35^. We observe a positive correlation in the gene-level phenotype across our screens (R=0.36-0.53 for 229 union gene set, **Figure S1F**). We find robust but incomplete correlation in the gene-level log2-fold-change in screens in serum-containing vs. serum-starved conditions (R=0.43 for 1800 genes in KO-Library 3, **Figure S1G**) and in our HepG2 serum-starved screens vs. previously screens in serum-starved HuH7 cells (R=0.50 for 203 genes in KO-Library 4, **Figure S1H**), suggesting that environmental conditions and cell line have moderate impacts on the genetic control of LDL-C uptake.

The screens reliably identify key known players in cholesterol uptake and metabolism, and flag intriguing novel candidate genes. Mutating known LDL-C uptake-implicated genes such as *LDLR* (the most robust hit), *SREBF2*^18,36^, *HNF4A*^37^, and *SCAP*^18,36^ impairs LDL uptake, while mutating *MYLIP*^28^ and cholesterol biosynthesis and processing enzymes *ACAT2* and *SQLE*^36,38^ increase LDL uptake (**Figure 1F, Supplemental Table 2**). The vast majority of the significant genes have not been previously reported to be involved in cholesterol metabolism, suggesting this is a rich dataset to identify novel genes and pathways involved in LDL-C uptake. To address the accuracy of this screening data, we cloned 20 individual gRNAs targeting genes found to impact LDL-C uptake and performed fluorescent LDL-C uptake assays on all 20 knockouts in triplicate. We find strong correlation between changes in LDL-C uptake in our CRISPR screen and individual validation (R^2^=0.69, **Figure 1G, Figure S1I**), confirming the accuracy of the CRISPR screening results.

### CRISPR activation screening reveals tunable regulators of LDL-C uptake

CRISPR-Cas9 knockout screens reveal genes that are required for proper LDL-C uptake. To determine which of these genes alter LDL-C uptake upon overexpression, we performed systematic upregulation of 200 LDL-C uptake-altering genes using CRISPR activation (CRISPRa) screening. To maximize the potency of CRISPR-based gene upregulation, we combined several approaches that have been shown to improve CRISPRa potency. We employed the SunTag system, in which dCas9 is appended with a string of 10 antigen peptides (dCas9-10xGcn4), allowing site-specific recruitment of 10 copies of a single chain antibody (ScFvGcn4) attached to transcriptional activation domains^39^. We fused ScFvGcn4 with an optimized combination of three human activation domains (SBNO1-NFE2L1-KRT40) found to provide stronger gene upregulation than the previously used VP64 peptide (Xu, Akinci *et al*, manuscript in preparation). Finally, we recruited two additional transactivators, p65 and HSF1, to the target site through fusion with a protein, MS2 coat protein (MCP), which binds to an MS2 hairpin sequence embedded in the gRNA^40^ (**Figure S2A**). We show that this combined CRISPRa platform leads to significantly stronger gene activation than the individual SunTag and MCP components (**Figure S2B**).

In addition to maximizing the potency of the CRISPRa platform, we sought to characterize the reliability of CRISPRa screening. Previous high-throughput CRISPRa studies have found CRISPRa to have more variable efficiency than CRISPRi^41,42^. These efforts have led to guidelines to select gRNAs predicted to be most active at inducing expression of a given gene^41,42^; however, these studies still find CRISPRa efficiency to be less predictable than that of CRISPRi or Cas9-nuclease. To measure the variable efficiency of gRNAs in CRISPRa screening, we have developed and employed the CRISPRa Outcome and Phenotype (COP) platform.

To carry out COP screening, we designed a high-throughput pooled lentiviral CRISPRa library in which each promoter-targeted gRNA has two targets: a target in an adjacent barcoded reporter construct containing a 205-nt proximal promoter sequence and a native genomic target in the promoter of a target gene (**Figure 2A**). This construct also expresses MCP-p65-HSF1. For each of 200 target genes, we cloned eight on-target gRNAs predicted to have high activity^42^ as well as two non-targeting control gRNAs. After transduction of HepG2 cells stably expressing dCas9-10xGcn4 and ScFvGcn4-SBNO1-NFE2L1-KRT40 in three biological replicates, we performed fluorescent LDL uptake screening, sorting the cells with the top and bottom 30% of LDL-C uptake to assess the degree to which each gRNA alters LDL-C uptake through activation of its native target gene. Additionally, we collected RNA to assess the magnitude of promoter reporter activation upon gRNA treatment through the transcribed barcode (**Figure S2A**).

**Figure 2:**
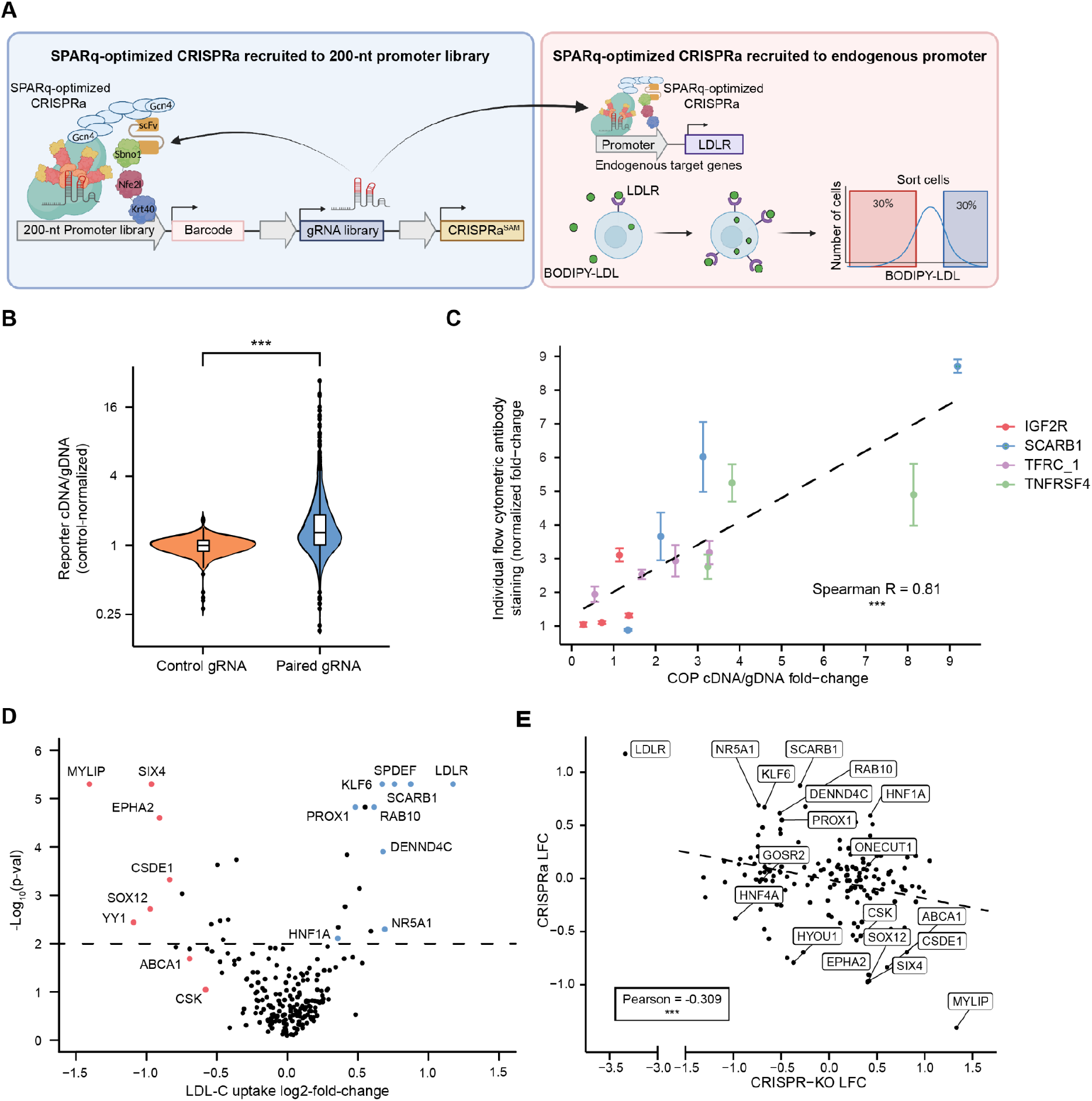
CRISPR activation screening reveals tunable regulators of LDL-C uptake. A: Schematic of CRISPRa Outcome and Phenotype (COP) screening approach. A gRNA is cloned into a lentiviral vector paired with a 205-nt region of its target promoter driving expression of a barcoded reporter transcript (CD5FLAG2). HepG2 cells expressing all CRISPRa components (see text) are treated with this gRNA library followed by selection. CRISPRa outcome analysis is performed through collection of genomic DNA (gDNA) and RNA from infected cells followed by library prep to assess reporter activation as the ratio of barcode representation in cDNA vs. gDNA. CRISPRa phenotype analysis is performed through fluorescent LDL-C uptake flow cytometric screening followed by MAGECK analysis to identify gRNAs and genes that alter LDL-C uptake. B: Reporter cDNA:gDNA ratio of all promoter-paired gRNAs vs. control gRNAs, normalized by the average cDNA:gDNA ratio of control gRNAs for each gene. Mann-Whitney U-test. P-values are represented by asterisks (*p<0.05, **p≤0.01, ***p≤0.001). C: Spearman correlation analysis of individual flow cytometric cell surface antibody staining and reporter cDNA:gDNA ratio. n=2-4. D: Volcano plot showing the MAGECK LDL-C uptake log2-fold-change (x-axis) and minimum MAGECK −Log10-p-value (y-axis) for the 200 genes in the CRISPRa library. Several significant genes are listed. E: Comparison of LDL-C uptake log2-fold-change for 200 genes targeted by CRISPR-KO (x-axis) and CRISPRa (y-axis).

Analysis of the promoter reporter found that the majority of gRNAs are active in CRISPRa. On-target gRNAs induced significantly higher reporter activity than control gRNAs (p<0.0001, measured as the ratio of cDNA:gDNA representation for all barcodes associated with a given gRNA), and 75% of on-target gRNAs produced stronger reporter activity than the median promoter-matched control gRNA (**Figure 2B, Supplemental Table 3**). We found that gRNA cleavage efficiency predictors such as Deep SpCas9 and CRISPick^43,44^, gRNA binding strand, as well as gRNA position with respect to the transcription start site showed significant correlations with CRISPRa activity (**Figure S2C**); however, a linear model incorporating such features showed weak predictive value of gRNA activity.

We found that reporter activity provides an accurate metric of CRISPRa activity. We evaluated the endogenous activation strength of 15 gRNAs targeting four cell surface proteins that were also evaluated in the COP screen, using flow cytometric antibody staining to measure CRISPRa potency. All four cell surface proteins were robustly activated by at least one tested gRNA, and the correlation between COP-measured activation and endogenous activation was high (Spearman R=0.81, p<0.001, **Figure 2C**, **Figure S2D**). Additionally, reporter activity was significantly, albeit incompletely, correlated with phenotypic impact on LDL uptake for on-target gRNAs targeting the 10 genes with strongest phenotypes (R=0.38, p<0.001, **Figure S2F**). As the vast majority of gRNAs showed CRISPRa activity, we found that MAGECK-RRA^45^ analysis of the LDL-C uptake screening data including all on-target gRNAs provided more statistical power than alternative analytic approaches that exclusively focused on active gRNAs.

We found that gene upregulation robustly alters LDL-C uptake. The ratio of gRNA representation in top vs. bottom-sorted cells was highly replicate-consistent (R=0.43-0.69, **Figure S2E**). Upregulation of 32 genes was found to significantly alter LDL-C uptake (FDR<0.1), with *LDLR* upregulation most robustly increasing and *MYLIP* upregulation most robustly decreasing uptake (**Figure 2D, Supplemental Table 4**). Comparing the effects of gene knockout vs. upregulation, we found highly significant anti-correlation (R=-0.31, **Figure 2E**), revealing that, on the whole, LDL-C uptake is a tunable process controlled by a set of core regulators.

### Gene module analysis elucidates LDL-C uptake-altering mechanisms

To gain insight into pathways that drive LDL uptake, we have employed co-essentiality analysis to group genes into functionally coherent modules. Databases of curated experimental data such as gene ontology (GO) and protein-protein interaction (STRING)^46–50^ rely on prior annotation of gene function, limiting their ability to classify poorly characterized genes. Co-essentiality analysis provides an unsupervised, hypothesis-free gene network based on the premise that genes with similar cellular functions will have similar fitness impacts in the same subsets of cell lines. Taking advantage of vast genome-wide CRISPR cell essentiality screen datasets that profile >1,000 cancer cell lines (DepMap)^51^, genes with similar functions can be clustered into modules without a bias toward characterized genes^52–56^. We have combined features of published co-essentiality pipelines^55,56^, using ClusterOne^57^ (d=0.2) to group nearly twice as many genes into functional modules as GO and STRING. We then annotate each module using overlapping GO modules for interpretability, finding that 85% of modules have a significantly enriched GO term (**Supplemental Table 5**).

Employing co-essential module enrichment on the 490 significant LDL uptake-altering genes, we identify 33 modules significantly associated with LDL-C uptake (hypergeometric p<0.01, **Supplemental Table 5**). The most significantly associated modules contain genes associated with vesicle-mediated transport, protein N-linked glycosylation, lipid biosynthesis, tRNA processing, and translation initiation (**Figure 3A, Figure S3A**).

**Figure 3:**
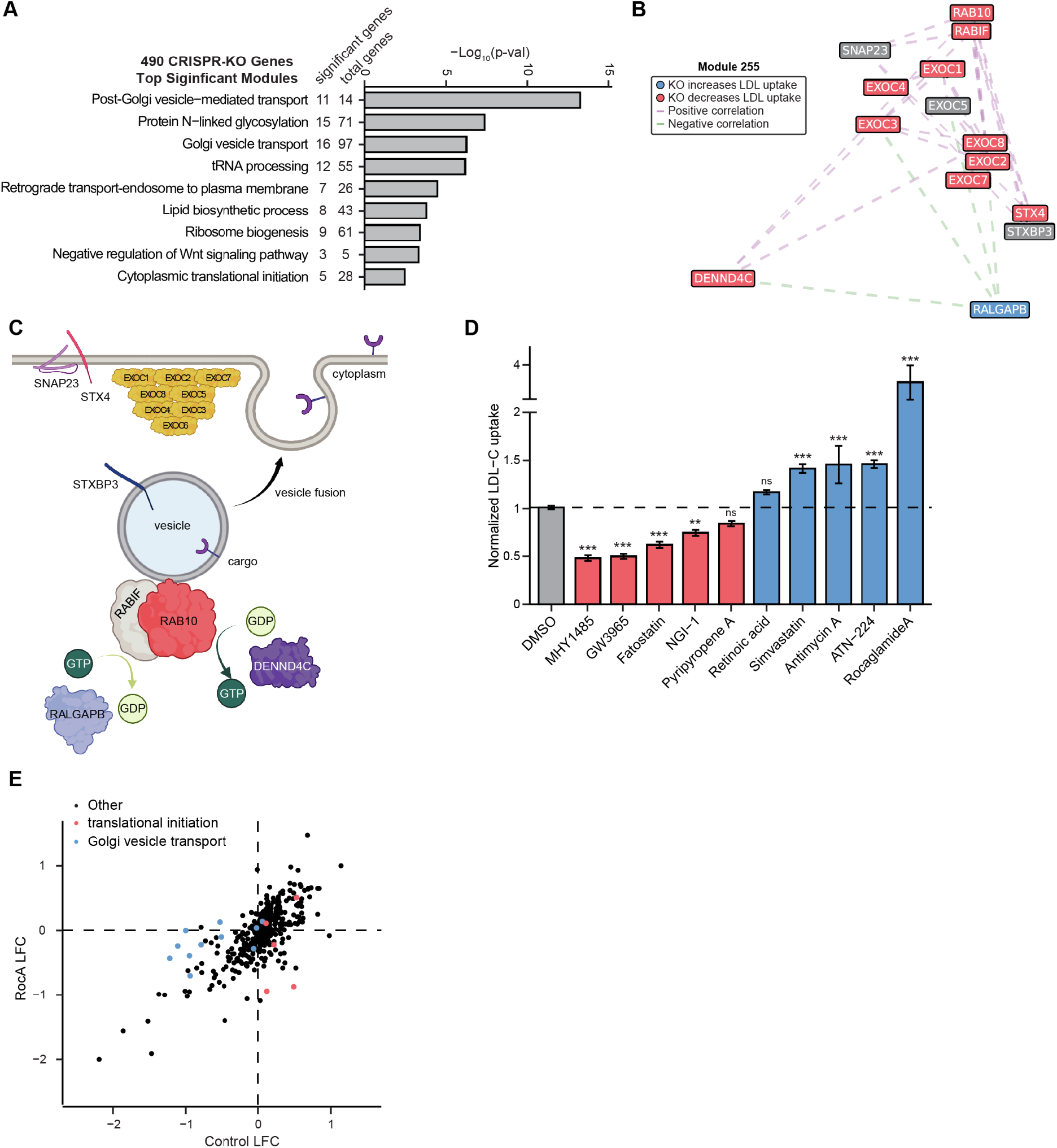
Gene module analysis elucidates LDL-C uptake-altering mechanisms. A: Co-essential modules with most significant enrichment of CRISPR-KO genes (hypergeometric p-value) labeled with the most significantly associated GO term for each module. B: tSNE plot based on PCA of module 255. Genes that decrease LDL-uptake when knocked out are labeled red, and genes that increase LDL-uptake when knocked out are labeled blue. Genes with pairwise Pearson correlation of >0.2 (purple) or <-0.2 (green) are connected with dashed lines. C: Module 255 contains RAB10 GTPase components including RAB10, the RAB10 holdase/chaperone RABIF, and the RAB10 GEF DENND4C and GAP RALGAPB. It also contains seven members of the exocyst complex that promotes vesicle exocytosis and three SNARE genes involved in vesicle fusion to the membrane. D: Mean control-normalized LDL-C uptake fold-change of HepG2 cells treated with DMSO (control) or one of 10 small molecules. Dunnett’s multiple comparisons test. n=6-26. P-values are represented by asterisks (*p<0.05, **p≤0.01, ***p≤0.001). E: Comparison of LDL-C uptake log2-fold-change for 522 genes targeted by CRISPR-KO in the presence of DMSO (x-axis) and Rocaglamide A (y-axis). Genes in co-essential modules annotated to be involved in translation initiation are shown in red and those involved in Golgi vesicle transport are shown in blue.

Most modules comprise a mix of genes previously annotated to be involved with a cellular function as well as previously unannotated genes^56^. To evaluate whether such unannotated genes contribute to the annotated cellular function, we explored module 146, whose most significant GO association is in protein N-linked glycosylation. 22 of 71 module members are annotated with this GO term, and 15 LDL uptake-altering genes are in this module, of which 7 are annotated with this GO term (**Supplemental Table 5**). We used a lectin flow cytometry assay in which we measure binding of two N-glycan-binding lectins, WGA and ECL, on the surface of HepG2 cells. We confirmed significantly reduced lectin binding upon knockout of two known, GO-annotated N-glycosylation genes that decrease LDL-C uptake, *OST4* and *STT3A* as well as upon treatment with the N-glycosylation inhibitor NGI-1 (**Figure S3B**). We tested an additional 5 genes in this module that are not annotated as N-glycosylation genes by GO, finding that knockout of 4 of these genes including two genes that decrease LDL-C uptake, *HYOU1* and *SLC39A7*, significantly reduce lectin binding. RNA-seq of HepG2 cells with *HYOU1* knockout reveals significant upregulation of co-essential modules involved in N-glycosylation (**Figure S3C, Supplemental Table 1**), suggesting compensatory upregulation of such genes. We note that reduced N-glycosylation induces pleiotropic defects in membrane trafficking and protein secretion^58^, and we conclude that dysfunction of many genes in module 146 decrease LDL-C uptake either directly or indirectly through altering the N-glycosylation of target proteins.

We also identify 13 significantly associated modules related to vesicle transport. The most significant, module 255, for which 11 of the 14 included genes significantly alter LDL-C uptake, contains RAB10 GTPase and exocytosis machinery (**Figure 3B-C**). Among RAB10-related genes, RAB10 is a GTPase implicated in numerous trafficking processes including Golgi-to-membrane and endosomal transport^59^, RABIF is a RAB10 holdase/chaperone that is required for RAB10 stability^60^, DENND4C is a RAB10 guanine exchange factor (GEF)^61^, and RALGAPB is a RAB10 GTPase activating protein (GAP). Module 255 also contains seven members of the exocyst complex, which is required for RAB10-mediated vesicle fusion with the plasma membrane^62^ as well as three SNARE complex proteins known to be involved in exocytosis^62^. Altogether, this cluster implicates exocytosis in LDL-C uptake, in line with previous findings^23,63^.

Beyond this exocytosis module, of the 321 total genes in these 13 vesicle-related modules, 53 are CRISPR screen hits. These modules highlight contributions of multiple distinct vesicle types in LDL-C uptake (**Figure S3D**)^64^, including those involved in the multivesicular body (module 514), the WASH and CCC endosome sorting complexes^65^ (modules 11, 277), endosome recycling (modules 476, 165), vesicle ATPase acidification (modules 229, 514), and ER-Golgi trafficking (modules 98, 372, 461), in addition to the exocyst complex and polarized transport machinery (modules 255, 69)^23,66^. Thus, co-essential clustering allows dissection of the multiple roles of vesicle trafficking in LDL-C uptake.

To assess the roles of co-essential modules on LDL-C uptake, we identified 10 small molecules that modulate the function of significant genes and pathways from our screens. Eight of the 10 molecules significantly alter LDL-C uptake in the direction expected based on the CRISPR screening data (**Figure 3D**). As a control, we tested 14 small molecules chosen for their effect on DNA repair^67^ and thus not based on the CRISPR screening data, finding that only 6 of 14 significantly alter LDL-C uptake (**Figure S3E**).

Reasoning that gene knockouts may be redundant or synergistic with small molecules that target the same cellular pathway^68^, we performed CRISPR screening using the 522-gene gRNA library of LDL-C uptake-altering genes in the presence of 8 small molecules as well as a DMSO control, each in three replicates. We found robust evidence of gene-small molecule interactions. For example, while *EIF4A1* knockout robustly increased LDL-C uptake in cells treated with DMSO or 7 unrelated small molecules, it decreased LDL-C uptake in cells treated with the EIF4A1 inhibitor Rocaglamide A, presumably due to redundancy with the drug (**Figure 3E, Supplemental Table 6**). Similarly, the LDL-C uptake-increasing effects of knockout of lipid biosynthesis pathway members *LSS* and *SQLE* were less pronounced in cells receiving the HMGCR inhibitor Simvastatin, presumably because Simvastatin acts through the same LDL-C uptake-increasing mechanism (**Figure S3F, Supplemental Table 6**). These observations were supported by examination of co-essential modules with differential effects in the presence of each drug as compared to DMSO-treated cells. The “translational initiation” module containing EIF4A1 was strongly depleted in Rocaglamide A-treated cells, while the “lipid biosynthetic process” module was strongly depleted in Simvastatin-treated cells (**Figure 3E, Figure S3F, Supplemental Table 7**). In addition to these expected small molecule-gene module interactions, co-essentiality analysis identified additional modules that interact with each small molecule. The LDL uptake-inhibiting effects of the “Golgi vesicle transport” module were mitigated in Rocaglamide A-treated cells, as were the effects of the “vesicle-mediated transport” in Simvastatin-treated cells (**Figure 3E, Figure S3F, Supplemental Table 7**).

To identify cellular functions that contribute to LDL-C uptake through related mechanisms, we built a correlation matrix of the differential effects of each co-essential module on LDL-C uptake in the presence of the 8 small molecules. We find that certain co-essential modules cluster in their interaction with these small molecules. For example, modules related to vesicle transport and N-linked glycosylation cluster, as do modules related to translation, mitochondrial function, and lipid biosynthesis functions (**Figure S3G, Supplemental Table 8**). Thus, analysis of thousands of gene-small molecule interactions identifies sets of cellular functions that act in coordinated fashion to alter LDL-C uptake.

### Coding burden analysis reveals monogenic contributions to serum LDL-C levels

To measure the *in vivo* effects of damaging variants within the genes and pathways identified in cholesterol uptake CRISPR screens, we examined the effects of rare coding variants on serum LDL-C levels using exome sequencing data and patient LDL-C values from the UK Biobank (UKB)^69^. First, we analyzed the coding burden of rare putative loss-of-function (pLOF) and computationally-predicted damaging missense variants from a published analysis of 454,787 individuals^14^. Among the 459 CRISPR-KO significant genes with qualifying variants in this dataset, we identify 11 genes with an LDL-C burden at a 10% false discovery rate (FDR) using this approach (**Figure 4A**, **Figure S4A**, **Supplemental Table 9-10**). These genes include known LDL-C uptake regulators such as *LDLR, MYLIP*, and *HNF1A*, a recently reported LDL-C GWAS target *RRBP1*^70^ (carriers have decreased LDL-C), as well as previously unreported genes such as the transcription factor *ZEB1* (carriers have decreased LDL-C), a key driver of epithelial-mesenchymal transition^71^, and the RAB10 GEF *DENND4C* (carriers have increased LDL-C). To address whether CRISPR-KO genes are enriched in genes with LDL-C burden in this pLOF and damaging missense focused analysis, we performed 10,000 randomized trials of 459-gene sets in the exome sequencing cohort. The most common result was 0 significant genes (34.63% of trials), and only 2 of the 10,000 trials gave more than 11 significant genes (**Figure S4B**), supporting the idea that HepG2 LDL-C uptake CRISPR-KO screening is a valuable approach to improve identification of genes associated with serum LDL-C levels. It is important to note that KO-Library 1, which contributed to the identification of a subset of the CRISPR-KO significant genes, was designed to include genes adjacent to LDL-C GWAS-associated variants. Analysis of the 2,634 genes in KO-Library 1 finds that these genes are enriched in genes with LDL-C burden as compared to random sets of equal numbers of genes (p=0.0062). Thus, we cannot rule out that some enrichment in CRISPR-KO hits with coding burden could derive from this non-uniform starting gene set. Whole genome LDL-C burden analysis using this approach revealed 51 significant genes, of which 8 overlap with genes identified in the CRISPR-KO-restricted analysis (**Supplemental Table 9-10**).

**Figure 4:**
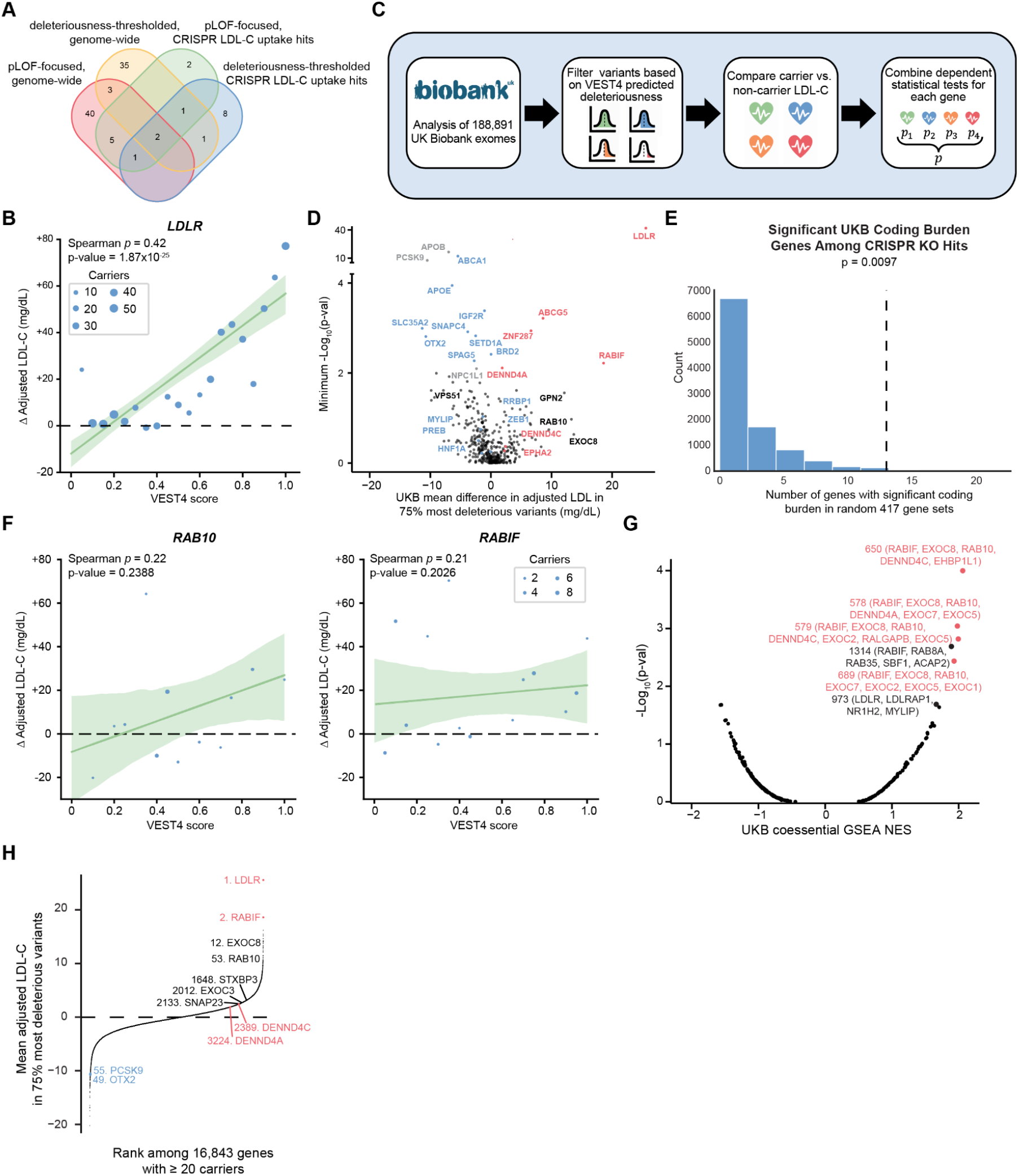
Coding burden analysis reveals contributions of genes and modules to serum LDL-C. A: Venn diagram showing genes with significant coding burden in the UKB cohort using two analysis methods. Both methods were employed genome-wide and restricted to CRISPR-KO-identified LDL-C uptake modulators. B: Correlation within UKB *LDLR* variant carriers between VEST4 score and change in adjusted LDL-C as compared to non-carriers. VEST4 scores are binned to the nearest 5%, the bubble sizes represent number of carriers within each bin, and a 95% confidence interval around the estimated linear regression line. C: Schematic of an approach to employ a collapsing burden model that accounts for variant deleteriousness to identify genes associated with altered serum LDL-C in the UKB exome cohort. D: Volcano plot showing the mean adjusted LDL-C difference (x-axis, mg/dL) and −Log10-p-value (y-axis) in the VEST4 burden analysis for 417 genes with significant effects on LDL-C uptake in the CRISPR-KO screen as well as 3 additional well-known LDL-C burden genes (gray). Genes significantly associated with increased (red) and decreased (blue) LDL-C levels in burden analyses are highlighted as well as selected non-significant genes (black). E. Bar chart displaying the number of genes significantly associated with serum LDL-C upon 10,000 random selections of 417-gene sets. Only 97/10,000 selections have at least as many significant genes (13) as when the 417 CRISPR-KO-significant genes are analyzed. F: Correlation within UKB *RAB10* and *RABIF* variant carriers between VEST4 score and change in adjusted LDL-C as compared to non-carriers. VEST4 scores are binned to the nearest 5%, the bubble sizes represent the number of carriers within each bin, and a 95% confidence interval around the estimated linear regression line. G: Hazard ratios and 95% confidence intervals from a Cox proportional hazard estimates of all rare missense and LOF variants’ effect on CAD development. H: Volcano plot showing the UKB LDL-C rank co-essential GSEA normalized enrichment score (NES) (x-axis) and −Log10-p-value (y-axis) for 241 co-essential modules with a ClusterOne threshold (d=0.8). The four significant modules (red) and several enriched non-significant modules are highlighted along with all genes included in those modules. I: Dot plot showing the rank in mean carrier adjusted LDL-C for the 75% most deleterious variants for 16,843 genes with 20 or more carriers. Select genes including those in the RAB10-associated co-essential module are highlighted with those significantly associated with hypercholesterolemia shown in red and hypocholesterolemia in blue.

The pLOF-focused coding burden analysis is restricted to variants which are predicted to be highly damaging^72^; however, such strict thresholding diminishes the ability to identify associations in genes which have low allele counts, either due to strong negative selection of damaging variants, or sequence length and context. In order to improve our sensitivity to detect such genes of interest which may have overlapping evidence from functional screening, we include a larger set of rare coding variants in a more inclusive burden analysis pipeline.

We evaluated 12 computational models that are designed to predict variant deleteriousness (*e.g*. CADD^73^, VEST4^74^) and evolutionary conservation scores (*e.g*. GERP^75^, PhyloP^76^). We assessed these models at two prediction tasks. First, we assess the predictive power of each model in classifying existing pathogenic and benign variant annotations from ClinVar^77^. We find that VEST4 outperforms other methods across classification thresholds (AUROC=0.96) for these variants (**Figure S4C**). Second, we assessed the correlation between deleteriousness scores from each of the 12 models and serum LDL-C levels for all 8,344 missense variants in 33 genes with significant coding burden^14^, adjusting serum LDL-C levels for age, sex, PRS, and statin use^5^ (adjusted LDL-C). We again found that VEST4 showed the highest mean correlation rank (**Figure S4D**) in this variant set. While the correlation between VEST4 score and adjusted LDL-C was modest and variable across these genes (median Spearman R = 0.077), the fact that correlations were significantly non-zero in 12/33 genes and robust in genes such as *LDLR, ABCA1*, and *ASGR1* (**Figure 4B**, **Figure S4E**) reinforced the idea that accounting for deleteriousness using VEST4 may improve the estimation of the impact of monogenic coding variants on LDL-C levels.

We thus assessed whether variants in the significant genes identified in our CRISPR-KO screens alter adjusted LDL-C in the UKB cohort by performing collapsing burden tests considering all variants in 188,891 individuals with four VEST4 score thresholds (top 25%, 50%, 75%, or 100% most deleterious, **Figure 4C**). Out of 417 CRISPR-KO significant genes with ≥ 10 carriers, we find 13 genes with LDL-C burden at a 10% FDR (**Figure 4A, 4D, Figure S4F, Supplemental Table 9, 11**), significantly more than are found using randomized 417-gene sets (**Figure 4E**, p=0.0097, CDF > 12 on 10,000 iterations). Interestingly, while the pLOF-focused and deleterious-thresholded burden tests identify similar numbers of significant genes, only four genes are identified in both analyses (*ABCA1, ABCG5, APOE*, and *LDLR*) (**Figure 4A**), revealing the complementary nature of these approaches. Genes uniquely identified in the deleterious-thresholded approach include the RAB10 holdase/chaperone *RABIF* (carriers have increased LDL-C), and the transcription factors *ZNF287* (carriers have increased LDL-C) and *OTX2* (carriers have decreased LDL-C). Whole genome LDL-C burden analysis using the deleterious-thresholded approach revealed 42 significant genes, of which 4 overlap with genes identified in the CRISPR-KO-restricted analysis (**Supplemental Table 9, 11**). VEST4 scores correlated reasonably well with UKB carrier LDL-C levels in genes with significant coding burden such as *OTX2* and *RABIF* as well as in other CRISPR-KO hit genes with robust impact on LDL-C levels such as *ACAT2, RAB10*, and *VPS51* (**Figure 4F**, **Figure S4G**), adding further evidence that deleterious variants in these genes lead to altered serum LDL-C levels.

We also examined rare variant associations with CAD among 469,828 UK Biobank exome sequencing participants in 21 genes. Using a Cox proportional-hazards regression, we find CAD to be significantly associated with LOF variants in *LDLR* (Hazard Ratio (HR) = 6.81 [10.16,4.56], p = 5.94×10^-21^) and *SPAG5* (HR = 2.08 [1.47, 2.96], p = 4.39×10^-5^), and missense variants in *LDLR* (HR = 1.28 [1.16,1.42], p = 2.29×10^-6^) (**Figure S4H, Supplemental Table 12**). These findings may be limited by statistical power given low numbers of rare LOF variant carriers, the number of CAD cases among variant carriers, or the broad spectrum of functional effects of missense variants. For example, in *PCSK9*, a gene where LOF variation is known to significantly decrease CAD risk, incidence at age 65 among LOF carriers is 1/70. As expected, there is a decreased odds ratio (0.2), but it is not significant (p = 0.13).

Among all genes with significance in the CRISPR-KO screens, there is a significant but weak negative correlation between their effect on LDL-C uptake and the effect of deleterious coding variants on serum LDL-C levels (Pearson r = −0.20, P<0.001, N=286, **Figure S4I**). Restricting to genes prioritized by both the CRISPR-KO LDL-C uptake phenotype and coding burden approaches, there is a significant and more robust negative correlation between their effect on LDL-C uptake and their effect on serum LDL-C levels (Pearson r = −0.79, P<0.01, N=10, **Figure S4J**). These results suggest that genes whose mutation decreases LDL-C uptake tend to lead to increased serum LDL-C, as would be expected.

In addition to gene-level burden analysis, we asked whether co-essential modules showed evidence of coherent association with serum LDL-C levels through gene set enrichment analysis (GSEA)^48^ using the ranked list of mean carrier serum LDL-C among variants in the top 75% of VEST4 deleteriousness for 16,843 genes with 20 or more qualifying variants. We evaluated co-essential modules at two ClusterOne thresholds (d=0.8 and d=0.2) to account for gene sets with high and low functional similarity. Among 241 d=0.8 modules that contain at least one significant gene in the CRISPR-KO screen, we found four significant modules, all associated with increased LDL-C levels, and comprising a total of 11 genes (**Figure 4G**, **Supplemental Table 13**). These genes can primarily be functionally designated as RAB10 GTPase components (*RAB10, RABIF, DENND4A, DENND4C, RALGAPB*) and members of the exocyst complex (*EXOC1, EXOC2, EXOC5, EXOC7, EXOC8*) (**Figure 3C**). Focused gene-level burden analysis on these 11 genes reveals that, among them, *DENND4A*, *DENND4C*, and *RABIF* have LDL-C burden (**Figure 4A**, **Supplemental Table 9**). Similarly, among 291 d=0.2 modules, the module with the strongest GSEA normalized enrichment score and the lowest p-value (p=0.007, not significant, **Figure S4K**, **Supplemental Table 14**) is module 255, which also comprises RAB10-related and exocyst pathway genes and scores as the most enriched module in the CRISPR-KO screen (**Figure 3B-C**).

These analyses indicate that RAB10 GTPase dysfunction contributes to hypercholesterolemia through impaired LDL-C uptake. Of the 16,849 genes assessed in the UKB exome cohort, deleterious variant carriers in *RABIF* have the second highest mean adjusted LDL-C (behind only *LDLR*), and carriers of *RAB10* variants have the 53rd highest mean adjusted LDL-C (**Figure 4H**). RAB10 GEFs *DENND4C* (2,389th of 16,849) and *DENND4A* (3,224th)^61^ are both significantly associated with serum LDL-C, albeit with weaker magnitude. Variant carriers in individual members of the exocyst complex such as *EXOC8* (12th) and *EXOC3* (2,012th) as well as exocytosis-promoting SNAREs *STXBP3* (1,649th) and *SNAP23* (2,134th)^62^ show elevated LDL-C, but heterogeneity within individual exocyst gene burden suggests that exocyst genes are not the main drivers of the LDL-C association of this module. Altogether, these analyses suggest that disruption of RAB10 vesicle-mediated processes, including exocytosis mediated by SNARE proteins and the exocyst complex, drives higher LDL-C levels in the human population.

### Disrupting the RAB10 vesicle transport pathway impedes LDLR membrane trafficking

We utilized individual CRISPR-KO experiments to study the role of RAB10 machinery in LDL-C uptake. We found through flow cytometry with an anti-LDLR antibody that HepG2 cells with CRISPR knockout of *RAB10* and *DENND4C* have significantly decreased surface LDLR levels (**Figure 5A**). To investigate the mechanisms driving this phenotype further, we repeated these knockouts in a Doxycycline-inducible LDLR-mCherry HepG2 cell line. Through timecourse fluorescence imaging, we found that LDLR is never efficiently trafficked to the cell membrane in *RAB10* knockout cells (**Figure 5B, Figure S5A**); instead, LDLR has a vesicular localization at all timepoints tested. In addition to altered localization, flow cytometric quantification revealed that *RAB10* and *DENND4C* knockout cells have significantly less LDLR-mCherry, indicating decreased LDLR stability (**Figure 5C**, **Figure S5B**). Finally, RNA-seq on *DENND4C* knockout HepG2 vs. control cells revealed enrichment in the expression of a co-essential module related to vesicle-mediated transport (**Figure 5D, Supplemental Table 1**); RNA-seq analysis of RAB10 knockout showed an elevated cellular stress signature and was thus excluded from analysis. It is of note that, in the co-essentiality dataset, *RAB10, RABIF*, and *DENND4C* cluster substantially more closely with exocyst complex members than with any other vesicle transport machinery, further suggesting that the primary role of *RAB10* in the cell is in facilitating exocytosis. Altogether, these data suggest that deficiencies in the RAB10 pathway lead to decreased membrane LDLR levels through decreased membrane trafficking and consequent decreased LDLR stability.

**Figure 5:**
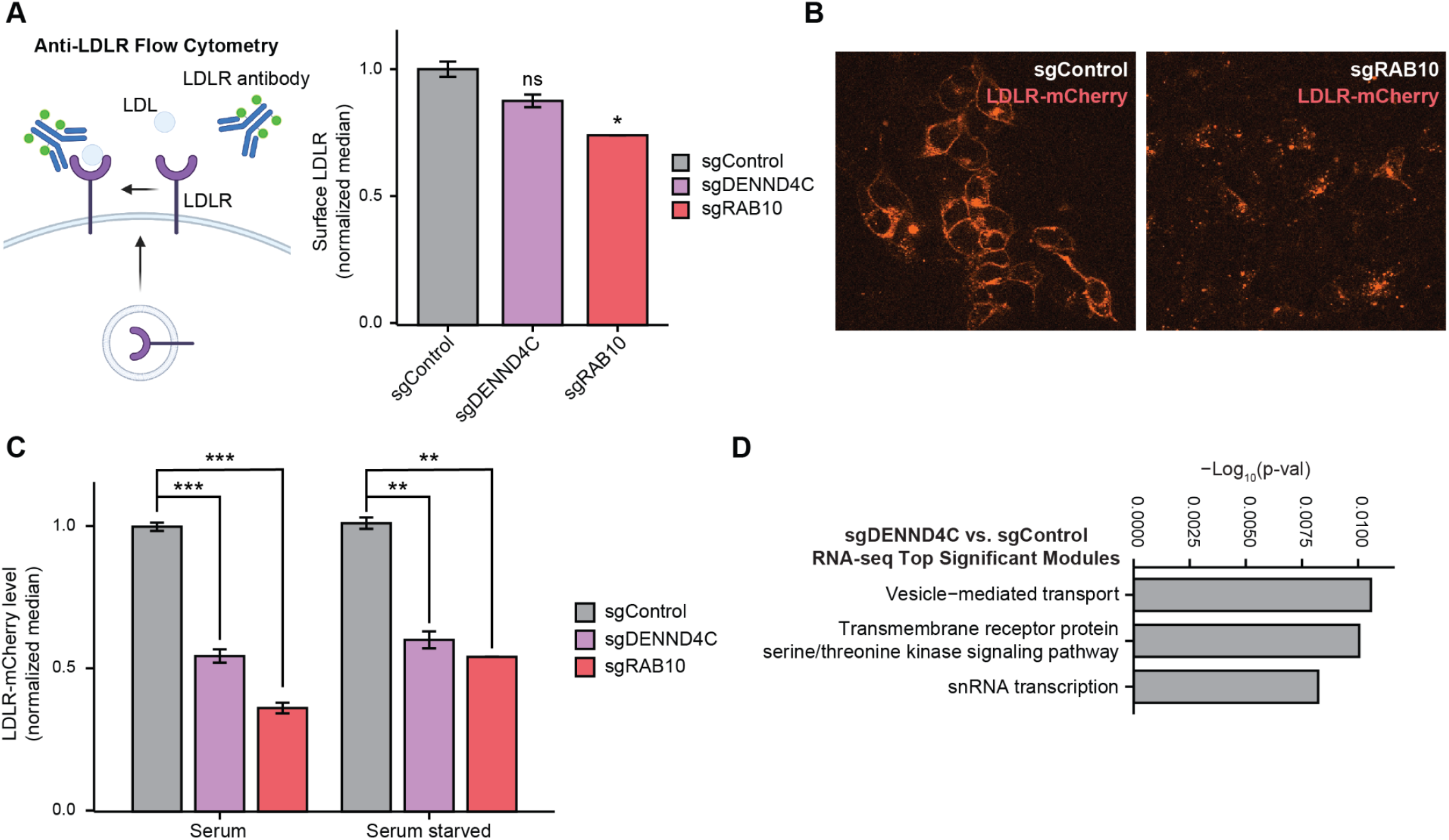
Disrupting the RAB10 vesicle transport pathway impedes LDLR membrane trafficking. A: Median anti-LDLR flow cytometric staining intensity for HepG2 cells with gene KO as indicated. Two-tailed t-test. n=2. B: Fluorescent microscopic images of HepG2 cells with inducible LDLR-mCherry expression and gene KO as indicated. LDLR-mCherry expression was induced 6 hours prior to imaging. C: Flow cytometric expression of LDLR-mCherry in HepG2 cells with inducible LDLR-mCherry expression and gene KO as indicated. LDLR-mCherry expression was induced 24 hours prior to measurement in the indicated media condition. n=2-4. Two-tailed t-test. P-values are represented by asterisks (*p<0.05, **p≤0.01, ***p≤0.001). D: Bar chart showing co-essential module enrichment among sgDENND4C RNA-seq upregulated genes.

### Mouse knockdown of Rabif, Otx2, and Csk alter serum cholesterol levels

To determine whether newly identified LDL-C uptake-altering genes affect organismal cholesterol levels, we asked whether the liver-specific knockdown of three such genes affects mouse serum cholesterol levels. We chose to study two genes with robust effects in our CRISPR-KO and CRISPRa screens as well as robust and statistically significant association with LDL-C in the UKB burden analysis, *RABIF* and *OTX2* (**Figure 4H**), and a third gene with robust *in vitro* phenotypes but without UKB burden, *CSK*. We confirmed that individual knockout of each gene significantly alters HepG2 LDL-C uptake and that ectopic expression of each gene’s open reading frame significantly alters LDL uptake in the opposite direction (**Figure 6A**). For *RABIF*, this was done through coordinated overexpression of *RABIF* and *RAB10*, given that these proteins seem to be coordinately required for proper function.

**Figure 6:**
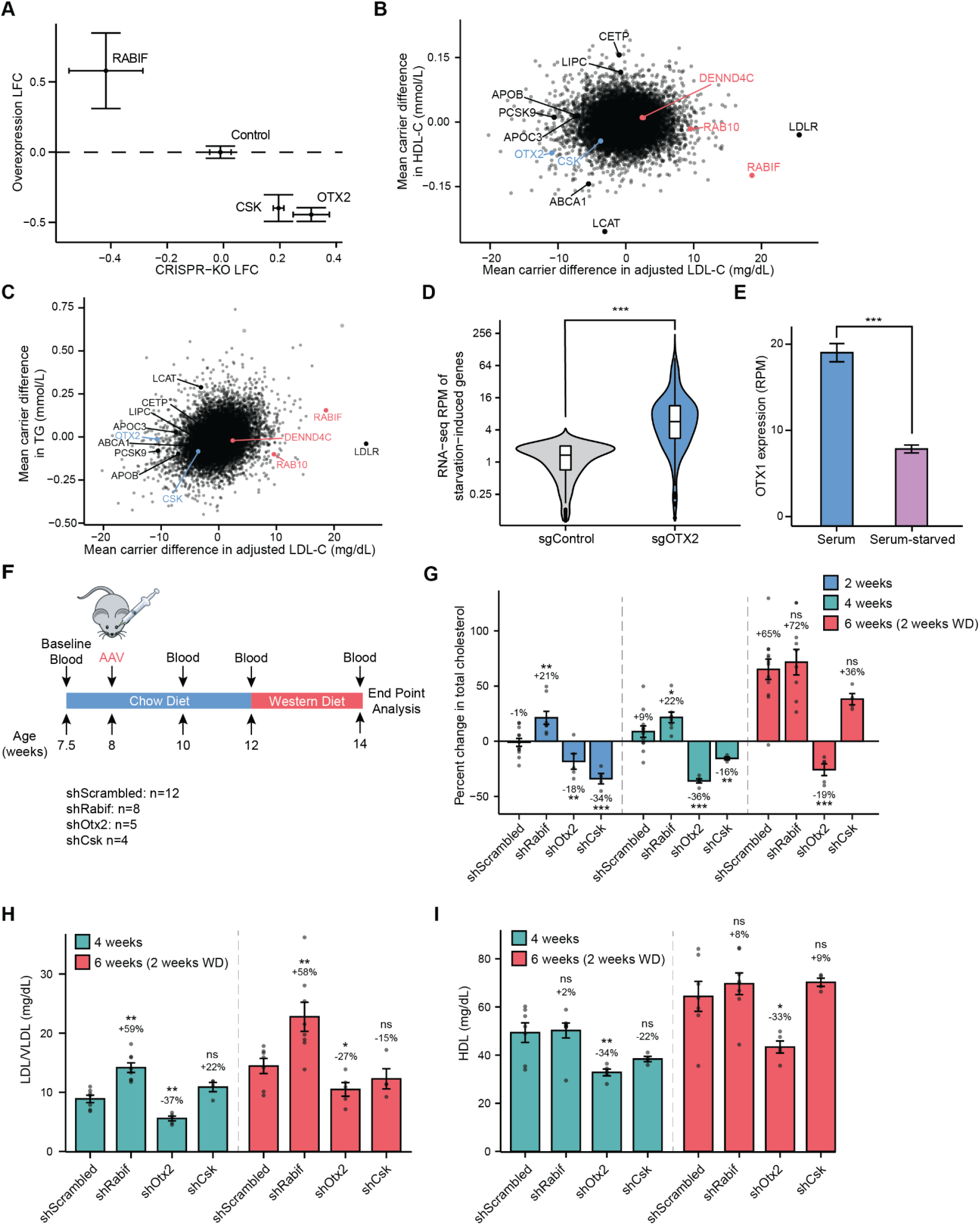
Mouse knockdown of Rabif, Csk, and Otx2 alter serum cholesterol levels. A: Comparison of the effects of CRISPR-Cas9 gene KO (x-axis) and ectopic ORF expression (y-axis) for CSK, OTX2, RABIF, and non-targeting control. Error bars represent standard error. B-C: Scatter plots showing UKB mean carrier adjusted LDL-C (x-axis) vs. mean carrier HDL-C (B, y-axis) and mean carrier triglycerides (C, y-axis) for all genes. Benchmark genes and novel genes are highlighted. D: HepG2 control or *OTX2* KO RNA-seq reads per million (RPM) for the 100 genes that are most robustly upregulated in HepG2 cells upon serum starvation. Paired Mann-Whitney U-test. E: RNA-seq RPM for *OTX1* in HepG2 cultured in serum-containing or serum-starved conditions. n=46. Two-tailed t-test. P-values are represented by asterisks (*p<0.05, **p≤0.01, ***p≤0.001). F: Schematic of mouse AAV shRNA experiment. Retro-orbital AAV8 shRNA injection is performed on 8-week old male mice followed by blood collection 2, 4, and 6 weeks post-injection. Western diet feeding begins 4 weeks post-injection, and endpoint analysis is performed 6 weeks post-injection. Cohort sizes for each treatment group are listed. G: Average percent change in total cholesterol from pre-treatment measurement in mice treated with the designated AAV8 shRNA at the designated timepoints. H: Average LDL/VLDL (mg/dL) in mice treated with the designated AAV8 shRNA at the designated timepoints. Average difference compared to shScrambled and p-value are noted. I: Average HDL (mg/dL) in mice treated with the designated AAV8 shRNA at the designated timepoints. Average difference compared to shScrambled and p-value are noted.

We also explored whether UKB carriers of deleterious coding variants in these genes show alterations in other serum lipids. *RABIF* carriers show low HDL-C and high triglyceride (TG) levels, although these trends do not rise to statistical significance (**Figure 6B-C, Figure S6A**). *OTX2* and *CSK* variant carriers show low HDL-C and middling TG, although also not statistically significant. Associations among these genes and Lipoprotein A and Apolipoprotein A levels are also predominantly non-significant.

We performed RNA-seq of HepG2 knockout cells to explore mechanisms underlying the LDL-C uptake-altering effects of *OTX2*. *OTX2* knockout induces robust upregulation of a serum starvation response gene program in non-starved HepG2 (**Figure 6D, Supplemental Table 1**). While *OTX2* expression is too low to reliably measure in HepG2 RNA-seq, its paralog *OTX1* is among the most highly downregulated genes upon HepG2 serum starvation (2.4-fold, p<0.001, **Figure 6E, Supplemental Table 1**). We note that *OTX1* also significantly increases LDL-C uptake in our CRISPR screen, suggesting these genes may play overlapping roles. We further performed co-essential GSEA on genes showing differential expression in *OTX2* knockout HepG2 cells to determine which gene modules show altered expression (**Supplemental Table 15**). Among the modules with the most upregulated expression in *OTX2* knockout vs. control HepG2 in serum-starved conditions are the “lipid metabolic process” module (module 660, p=0.002, FDR=0.44) and “cholesterol esterification” module (module 335, p=0.02, FDR=0.66), and the most downregulated module is the “triglyceride catabolic process module (module 324, p<0.001, FDR=0.14). Analysis of *OTX2* knockout vs. control HepG2 in serum conditions yielded a distinct set of altered modules, with the most enriched being the “protein transport” module (module 69, p<0.001, FDR=0.23), which contains the RAB10/Exocyst genes.

To further explore the gene regulatory activities of OTX genes, we analyzed published genome-wide Perturb-seq data in which RNA-seq profiling was performed after CRISPRi knockdown of 9,858 genes in the myelogenous leukemia cell line K562^78^. While knockdown of *OTX1* and *OTX2* were performed in this study, the *OTX2* knockdown experiment failed quality control because it did not produce a significant decrease in *OTX2* levels. To analyze the effect of *OTX1* knockdown, we performed co-essential GSEA to determine which gene modules show altered expression. The module with the most upregulated expression upon *OTX1* knockdown is the “regulation of cholesterol biosynthetic process” module (module 82, p=0.002, FDR=0.13, **Supplemental Table 16**). The most downregulated module (module 160, p<0.0001, FDR<0.0001) comprises genes involved in “SRP-dependent cotranslational protein targeting to membrane,” or the ER targeting of proteins destined for secretion or membrane deposition. While these data derive from a myelogenous leukemia cell line, which differ from the liver substantially in cholesterol processing, the upregulation of genes involved in regulating cholesterol biosynthesis is consistent with the HepG2 RNA-seq analysis and our data showing that dampened OTX function leads to increased LDL-C uptake and decreased serum LDL-C levels.

To begin to determine how OTX genes control these gene expression changes, we analyzed OTX1 ChIP-seq data from K562 cells generated by the ENCODE project. These data identify 10,690 OTX1 binding sites in the K562 genome (**Supplemental Table 16**). We used GREAT^79^ to identify 3,549 human genes with OTX1 promoter binding (+5 to −1 kb of the gene’s TSS). Transcription factor binding proximal to a gene does not necessarily equate to direct *cis*-regulation, so we asked whether OTX1 was more likely to bind proximal to genes whose expression was changed by *OTX1* knockdown. We found that the 10% of genes most upregulated upon *OTX1* knockdown were significantly more likely than other genes to have OTX1 promoter binding (36% vs. 31%, p<0.01), while genes downregulated upon *OTX1* knockdown had marginally decreased likelihood of OTX1 promoter binding (28% vs. 31%, p<0.05). Further, genes with OTX1 promoter binding have significantly higher expression upon *OTX1* knockdown than genes without such binding (median LFC 0.019 vs. 0.012, p<0.0001). These data provide supportive evidence that OTX1 acts more often as a transcriptional repressor than an activator, although OTX1 binding is likely to play more nuanced roles at individual binding sites. Due to the widespread binding of OTX1 across the K562 genome, it is difficult to assign causality to individual targets. OTX1 promoter binding is significantly enriched at promoters of genes in module 160 (51% vs. 31%, p<0.01) and not significantly different at module 82 promoters, suggesting that these module 160 genes involved in cotranslational ER targeting may represent key *cis*-regulatory targets of OTX1. Taken together, our data point to a role for *OTX* genes in regulating gene programs related to lipid metabolism and cholesterol biosynthesis across several cell states. *CSK* knockout induces the expression of co-essential modules related to cell motility and cytoskeleton organization (**Figure S6B**).

In order to evaluate the effects of these genes in mice, we performed AAV8 shRNA-mediated gene knockdown of all three genes using retro-orbital injection, which is known to yield efficient liver delivery^80–82^. We measured serum cholesterol levels prior to knockdown and then every other week for six weeks, with the first four weeks on standard chow and the final two on Western diet (WD) (**Figure 6F**). At the six week endpoint, we confirmed that shRNA treatment induced robust decreases in target transcript levels in liver (on average 88% decrease for *Rabif*, 60% decrease for *Csk*, *Otx2* transcripts were undetectably low, **Figure S6C**)

We observed rapid, robust, and durable shRNA-mediated alteration of serum cholesterol compared to pre-treatment baseline levels. *Rabif* knockdown increased total cholesterol levels by an average 21% as compared to pre-treatment levels within 2 weeks (N=8), significantly more than the 1% decrease observed in scrambled shRNA control mice (N=12, p<0.01). This serum cholesterol increase was maintained at 4 weeks (22% vs. 9% in shScrambled, p=0.05) and expanded at 6 weeks after 2 weeks on WD (72% vs. 65% in shScrambled, NS) (**Figure 6G**). The increased total cholesterol in sh*Rabif*-treated mice was primarily driven by changes in the VLDL/LDL fraction, which was 59% higher than in shScrambled-treated mice at week 4 and 58% higher at week 6 (N=8, p<0.01 at both timepoints, **Figure 6H**) with only minimal changes in HDL (N=8, NS, **Figure 6I**). *Rabif* knockdown also increased serum triglyceride levels by 13% as compared to control knockdown after 4 weeks, although this change was not significant (**Figure S6D**). Altogether, mouse liver-targeted knockdown of *Rabif* led to markedly increased serum LDL-C levels, in line with the higher serum LDL-C observed in individuals with deleterious *RABIF* variants and the decrease in liver cell LDL-C uptake after *RABIF* knockout.

On the other hand, knockdown of *Otx2* led to robustly decreased serum cholesterol as predicted from biobank and cell line analysis. *Otx2* knockdown decreased cholesterol levels by 18% and 36% at 2 and 4 weeks respectively (N=5, p=0.01 at 2 weeks, p<0.001 at 4 weeks) (**Figure 6J**). In comparison, liver-targeted siRNA-mediated knockdown and CRISPR-Cas9 mutation of *Pcsk9* have been shown to reduce total cholesterol by 30-40% in chow-fed wild-type mice^83,84^. *Csk* knockdown decreased total cholesterol levels compared to pre-treatment levels by an average of 34% and 16% at those timepoints (N=4, p<0.001 at 2 weeks, p<0.01 at 4 weeks). This pronounced decrease in cholesterol was maintained in sh*Otx2*-treated mice at week 6 after 2 weeks of WD (19% decrease, N=5, p<0.001) but not in sh*Csk*-treated mice (36% increase, N=4, NS) (**Figure 6G**). *Otx2* knockdown decreased both VLDL/LDL and HDL levels significantly as compared to control knockdown at 4 weeks (by 37% and 34%, N=5, p<0.01 for each) and 6 weeks (by 27% and 33%, p<0.05 for each) (**Figure 6H-I**), in line with human exome data, and *Otx2* knockdown also robustly decreased triglyceride levels as compared to control knockdown by 26% at 4 weeks (p<0.05, **Figure S6D**). *Csk* knockdown did not induce significant changes in separated VLDL/LDL or HDL levels at either 4 or 6 weeks (N=4), nor did it alter triglyceride levels (**Figure 6H-I**, **Figure S6D**).

Certain genetic mutations that lower circulating LDL-C levels, such as those in *APOB*, cause increased hepatic lipid and cholesterol levels, while other mutations such as those in *PCSK9* do not^85^. We assessed the effect of *Otx2* knockdown on liver total cholesterol and triglyceride levels. We found that mice with Otx2 knockdown show on average 18% reduction in liver total cholesterol (p<0.05) and 24% reduction in liver triglyceride levels (p<0.05, **Figure S6E**). These data demonstrate that *Otx2* reduces lipid and cholesterol content both in the blood and in the liver, suggesting that *Otx2* loss is unlikely to reduce circulating LDL-C through impairing VLDL secretion. We conclude that liver-targeted knockdown of *Otx2* leads to robust decrease in serum LDL-C levels and makes mice refractory to WD-induced increases in cholesterol level.

### LDL uptake-altering genes may underlie GWAS loci

GWAS-identified loci often have multiple candidate variants, which predominantly tend to be in non-coding regions, complicating their connection to CRISPR screen-identified genes. Nonetheless, we asked whether LDL uptake-altering genes we identified could potentially underlie loci associated with LDL-C in a recent trans-ancestry GWAS^15^. Examining the RAB10 co-essential module, we noted GWAS-identified loci significantly associated with LDL-C adjacent to *RAB10* and *DENND4C*^15^. Bayesian fine-mapping^86^ of the *RAB10* locus suggests a single likely causal variant (rs142787485, posterior inclusion probability (PIP)=0.92) in a conserved region of the 3’ UTR of *RAB10* (**Figure 7A)**. The minor allele of rs142787485, which is associated with lower LDL-C levels in carriers^15^, disrupts predicted binding sites of two microRNA species (**Figure S7A**)^87^, suggesting that the minor allele may possess increased *RAB10* transcript stability. To evaluate the impact of this variant, we cloned a GFP-RAB10 fusion protein expression construct containing the full RAB10 3’ UTR with rs142787485 major or minor allele. We found that, upon stable lentiviral transduction into HepG2 cells, GFP-RAB10 with a 3’ UTR with rs142787485^min^ led to 9% higher protein levels than rs142787485^maj^ (n=6, p<0.001, **Figure 7B-C**). Altogether, these data suggest that, in addition to rare coding variants in RAB10 pathway genes that associate with hypercholesterolemia, a common variant that increases RAB10 protein levels is associated with reduced LDL-C levels in the population. Fine mapping of the *DENND4C* locus yields 7 intronic variants with posterior inclusion probability (PIP) between 5-25% (**Figure 7D)**, complicating evaluation of regulatory mechanisms.

**Figure 7:**
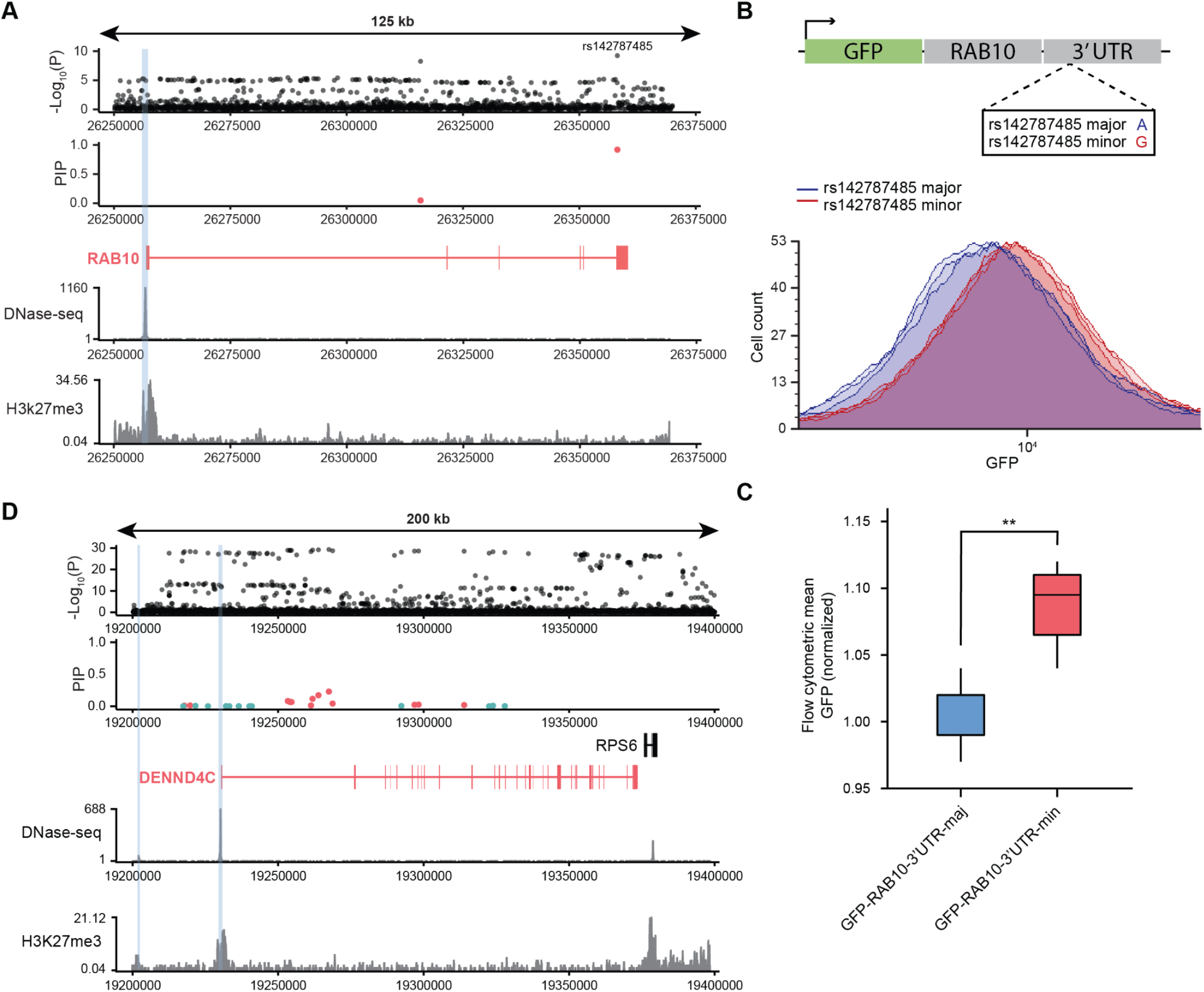
LDL uptake-altering genes may underlie GWAS loci. A: Fine-mapping of the GWAS signal at the *RAB10* locus (black: LDL-GWAS variants, red: 95% confidence set, teal: 99% confidence set). Signal tracks represent chromatin accessibility from DNase-seq or H3K27me3 histone modifications from ChIP-seq. Blue bars represent ABC predicted enhancer connections with *RAB10*. B: Flow cytometric analysis of HepG2 cells with expression of a GFP-RAB10 fusion protein expression construct containing the full RAB10 3’ UTR with rs142787485 major (blue) or minor (red) allele. C: GFP-RAB10 3’ UTR mean GFP flow cytometry. n=6. Two tailed t-test. P-values are represented by asterisks (*p<0.05, **p≤0.01, ***p≤0.001). D: Fine-mapping of the GWAS signal at the *DENND4C* locus (black: LDL-GWAS variants, red: 95% confidence set, teal: 99% confidence set). Signal tracks represent chromatin accessibility from DNase-seq or H3K27me3 histone modifications from ChIP-seq. Blue bars represent ABC predicted enhancer connections with *DENND4C*.

Other robustly LDL-uptake altering genes that are not classically recognized cholesterol modulators but which are nearby to LDL-C GWAS loci include *CSK*, *ARID1A*, and *RRBP1* (**Figure S7B-D**). In particular, the *ARID1A* locus contains a single credible fine-mapped variant in the *ARID1A* promoter (rs114165349, PIP=0.99), which is predicted to regulate its expression by the ABC model (**Figure S7B**). ARID1A is a core epigenetic regulator whose loss in mouse liver leads to pleiotropic changes in gene expression, cellular function, and lipid metabolism, including elevated cholesterol^88^. Additionally, the GWAS signal at the *RRBP1* locus can be localized to a single fine-mapped variant (rs2618566, PIP=1.00) in a distal regulatory region that is a liver eQTL^70^ (**Figure S7C**). Our exome burden analysis identifies significantly decreased LDL-C in carriers of rare *ARID1A* and *RRBP1* coding variants (**Figure S7E**), and mouse *RRBP1* knockdown has been shown to disrupt lipid homeostasis and lead to lower circulating LDL-C levels^89^. Nonetheless, further experiments will be required to determine which variants surrounding these genes are causal and whether these predominantly non-coding GWAS variants do indeed alter expression or function of these LDL uptake-altering genes.

## Discussion

LDL-cholesterol has long been recognized as a risk factor for CAD^90^, and genetic contributions to cholesterol levels have been known for over 100 years^91^. Nonetheless, genetic contributions to LDL-C levels remain incompletely understood, as evidenced by the hundreds of GWAS-associated loci without nearby known LDL-C regulatory genes. We have employed a pipeline combining human cell line CRISPR screening, co-essentiality-based gene module analysis, human genetic association, and mouse knockdown to expand the understanding of genes and cellular mechanisms whose dysfunction alters serum LDL-C levels. Overall, combining our newly identified burden genes with those from published genome-wide analyses^11–14^, we expand the list of genes significantly associated with LDL-C levels to 101 genes (**Supplemental Table 9**). Of these genes, 21 function at least partially through altering liver cellular LDL-C uptake. This gene set has already proven a valuable resource to understand pathways and mechanisms altering cholesterol metabolism, to evaluate therapeutic targets, and to connect GWAS-identified variants with target genes.

Combining human genetic association and cellular genetic screening compensates for shortfalls in each individual approach. While genetic association cohort sizes are ever increasing^14^, the rarity of deleterious coding variants and the genetic and environmental heterogeneity among carriers complicates the identification of disease-associated genes from human cohorts alone. CRISPR screening, on the other hand, allows highly scalable and well controlled loss-of-function screening; however, variability in cellular LDL-C uptake is not a perfect surrogate of serum LDL-C levels. Combining these approaches allows us to focus coding variant examination on a subset of genes with heightened probability of impacting LDL-C levels in the population. We find that focusing coding burden analysis specifically on genes with *in vitro* LDL-C uptake phenotypes increases power to identify significant associations. This is not simply the result of decreasing the number of tested hypotheses– LDL-C uptake-impairing genes are highly enriched in genes with LDL-C burden (**Figure 4E, Figure S4E**), although we cannot rule that some portion of this enrichment derives from the enriched starting gene set we employed in KO-Library 1.

These previously unappreciated burden genes often show robust effects on LDL-C levels in deleterious variant carriers, as with *RABIF* and *RAB10* for which carriers have the 2nd and 53rd highest mean LDL-C among 16,849 evaluated genes and *OTX2* for which carriers have the 49th lowest mean LDL-C, on par with *PCSK9* (**Figure 4G**). We validate the impacts of dampened expression of two such genes in mice, showing effects on LDL-C levels that are similar in magnitude to those of known monogenic hyper/hypocholesterolemia genes.

These genes have likely eluded prior identification due to the relative rarity of deleterious variant carriers. Classical hypercholesterolemia and hypocholesterolemia genes have been identified through characterization of human loss-of-function alleles and knockout mice^16^. However, germline knockouts of *Otx2, Rab10, Dennd4c*, along with many other genes we implicate in LDL-C uptake, are embryonic lethal^92–95^, and humans with bi-allelic disruption of these genes have not been reported. Additionally, genes such as *RAB10* and *RABIF* are small (200 and 123 amino acids respectively), making them inherently less likely to accumulate mutations. The techniques that we utilize, including cell type-specific CRISPR screening and analysis of mono-allelic partial loss-of-function phenotypes from exome biobank data, enable the identification of disease-relevant genes that are not easily identified using organism-wide loss-of-function-based approaches. Moreover, organ-specific therapeutic delivery approaches such as GalNAc-conjugated siRNA^96^ provide platforms for targeted manipulation of such genes.

Additionally, we find that co-essentiality provides a powerful approach to grouping genes into functional modules. Co-essentiality annotates a larger number of genes than GO and PPI database analysis^56^ and, because it is based on functional impacts of gene KO, is well-calibrated for pathway analysis of CRISPR screening data. Of note, through the combined deployment of CRISPR screening, co-essentiality analysis, and exome burden analysis, we define a previously unappreciated pathway contributing to genetic propensity toward hypercholesterolemia. We find that the RAB10-driven exocytosis pathway shows significant LDL-C burden.

RAB10 is a multi-functional GTPase that has been implicated in a diverse array of vesicle-mediated processes including endocytosis and exocytosis^59,63^; however, the co-essentiality data suggests stronger shared function with the exocyst complex than other vesicle-mediated processes, and we show that RAB10 pathway mutants show profoundly decreased LDLR membrane trafficking. The exocyst complex has been previously implicated in polarized transport of LDLR to basolateral membranes^66^, and mutations in exocyst genes impair LDLR uptake while increasing transferrin uptake^23^. Exocytosis of LDLR is vital for initial plasma membrane delivery as well as for endosomal recycling to the membrane, and thus it is likely that defects in the RAB10/exocyst pathway act at multiple stages in LDLR membrane trafficking. In our timecourse microscopy data, we are unable to detect substantial surface LDLR expression in RAB10-mutant cells at its earliest detectable timepoint, suggesting that impaired initial membrane trafficking indeed contributes to the phenotype. Recent work has also shown that RAB10-deficient cells show defective LDLR endosomal recycling as well^63^. Additionally, recent work has shown that RAB10 and exocyst knockout impairs LDL-C uptake in LDLR knockout cells^23,63^, and RAB10 has been implicated in lipophagy^97^, suggesting that this pathway’s roles in lipid uptake and trafficking are multifaceted, going beyond LDLR membrane deposition. These additional roles of RAB10 may explain why the LDL-C exome burden within the enriched module clusters most strongly with *RAB10*, *RABIF*, and its GEFs *DENND4C* and *DENND4A* and less so with exocyst genes.

The RAB10 pathway regulates LDL-C uptake quantitatively. Knockout of *RAB10*, its stability-promoting chaperone *RABIF*, and the RAB10 GEF *DENND4C* decrease LDL-C uptake while knockout of the RAB10 GAP *RALGAPB* increases uptake. Conversely, our CRISPRa data indicates that upregulation of *RAB10* and *DENND4C* robustly increases LDL-C uptake. Our analysis of the UKB exome cohort also primarily detects quantitative trait regulators, as the deleterious variant-associated effects we measure are mono-allelic in the affected individuals. Finally, a recent GWAS identifies common variants nearby *RAB10* and *DENND4C* that are associated with LDL-C levels, and we validate that the RAB10 3’ UTR variant that is associated with decreased serum LDL-C levels increases RAB10 protein levels by 9% in liver cells. Thus, our work identifies common and rare variation in LDLR exocytosis as a contributor to serum LDL-C levels, adding another point of regulation beyond the cellular decision to recycle or degrade LDLR which is influenced by PCSK9 and MYLIP. The RAB10/exocytosis pathway is not transcriptionally regulated in response to starvation in HepG2 (**Supplemental Table 1**) or in mouse hepatocytes^98^, but it remains to be seen whether this pathway acts as a feedback regulator of LDLR levels under other conditions and whether it could be activated for therapeutic benefit.

Further, we show that decreased function of *OTX2* robustly lowers LDL-C levels in human cohorts and mice through increased LDL-C uptake. *OTX2* emerges as a compelling candidate for further study given the hypocholesterolemic effect of its disruption. However, unlike genes with similar hypocholesterolemic effects such as *PCSK9* and *ANGPTL3* where human loss-of-function alleles are clearly associated with decreased CAD^99,100^, deleterious *OTX2* alleles are too rare to ascertain effects on CAD. *Otx2* is essential for gastrulation and early brain and eye development^92^; its roles outside of the nervous system are poorly understood. *OTX2* RNA expression is undetectably low in human and mouse liver tissue and in HepG2 cells, yet its downregulation in HepG2 and mouse liver yields a strong transcriptional and cholesterol phenotype. Knockout of the orthologous and partially functionally redundant *OTX1* gene^101^ in HepG2 yields similar increase in HepG2 LDL-C uptake, and individuals with deleterious *OTX1* variants have lower LDL-C levels, albeit not as robustly as for *OTX2*. Whether dual inhibition of *OTX1* and *OTX2* more dramatically lowers LDL-C levels and whether hepatic *OTX2* inhibition has side effects beyond its effects on serum lipids are compelling areas for further research.

In this work, we focus on genetic control of liver cell LDL-C uptake, which is but one of many points of control of serum LDL-C levels. As such, we posit that many additional genes are likely to be *bona fide* LDL-C regulators that could be validated through expanded exome sequencing cohorts combined with cellular assays for distinct phenotypes known to contribute to organismal LDL-C levels. As an example, a number of genes in the lipid biosynthesis co-essential module associate with substantially lower LDL-C levels (**Figure S7F, Supplemental Table 11**), yet only the statin target HMGCR rises to statistical significance. Likewise, carriers of deleterious variants in the pancreatic colipase gene (*CLPS*) have robustly decreased LDL-C levels that do not reach statistical significance (**Figure S7F, Supplemental Table 11**). The drug orlistat inhibits the effects of CLPS in the intestine, and orlistat use has been shown to associate significantly with decreased LDL-C levels^102^. In addition to expanding the range of phenotypic assays, it will also be important to evaluate exome sequencing cohorts from diverse ancestries to improve understanding of drivers of cholesterol metabolism across genetic backgrounds.

Altogether, the integrated approach combining cellular genetic screening, biobank genetic association, and gene network analysis defined here should provide a roadmap to guide efforts to dissect complex trait genetics.

## STAR Methods

### Resource availability

#### Lead contact

Further information and requests for resources and reagents should be directed to and will be fulfilled by the lead contact, Richard I. Sherwood

#### Materials availability

Plasmids generated in this study will be deposited to Addgene to be publicly available as of the date of publication.

#### Data and code availability

- RNA-seq and CRISPR screening data will be deposited at GEO and will be publicly available as of the date of publication. Accession numbers will be listed in the key resources table.
- All original code will be deposited at https://github.com/j-fife/hamilton-et-al and will be publicly available as of the date of publication. DOIs will be listed in the key resources table.
- Any additional information required to reanalyze the data reported in this paper is available from the lead contact upon request

### Experimental model and subject details

#### Cell lines

HepG2 hepatocellular carcinoma cells

#### Animal models

C57BL/6 mice

### Method Details

#### CRISPR-KO Screening

For KO-Library 1, four gRNAs were designed targeting each of the 2,634 genes within 1 Mb of a lead variant from the MVP multiethnic GWAS cohort^3^. gRNAs were selected from the Brunello, TKOv3, and CROATAN libraries^44,103,104^ to maximize inDelphi-predicted frameshift fraction^29^. 200 non-targeting control gRNAs were chosen from the Brunello library. KO-Library 2 included the same four gene-targeting gRNAs from KO-Library 1 for 360 genes that showed strong phenotypic effects as well as the same non-targeting controls. KO-Library 3 contained the 4 Brunello gRNAs for 1,803 transcription factor genes^32,33^ as well as Brunello non-targeting controls. KO-Library 4 included four Brunello gRNAs targeting 522 genes with significant or near-significant effects on LDL-C uptake from the three previous screens as well as from a recently published genome-wide LDL-C uptake CRISPR-Cas9 screen performed in Huh7 cells^23^ and also included Brunello non-targeting controls.

All oligonucleotide libraries (**Supplemental Tables 17-20**) were ordered from Twist Biosciences or Agilent Technologies in the following sequence format:

CTTGTGGAAAGGACGAAACACCG [19-20-bp protospacer—remove initial G for any 20-bp protospacer with one natively] GTTTAAGAGCTATGCTGGAAACAGCATAGC

Libraries were amplified by PCR using Q5UltraII mastermix (NEB) using the following primers: gRNA_60bp_fw TAACTTGAAAGTATTTCGATTTCTTGGCTTTATATATCTTGTGGAAAGGACGAAACACCG gRNA_60bp_rv GTTGATAACGGACTAGCCTTATTTAAACTTGCTATGCTGTTTCCAGCATAGCTCTTAAAC gRNA libraries were cloned into the lentiCRISPR-v2-FE-PuroR backbone (derived from Addgene plasmid #52961^105^ by replacing the gRNA hairpin with the FE gRNA hairpin^106^) through BsmBI vector digest and NEBuilder HiFi DNA assembly, ensuring >100-fold representation of each gRNA.

HepG2 Cas9NG-P2A-mCherry cells were seeded at a density of 4×10^4^ cells/cm^2^ on 15 cm plates in four biological replicates with 8 μg/mL of polybrene. The cells were transduced with lentivirus using Trans-IT Lenti (Mirus Bio) at a multiplicity of infection (MOI) of 0.3-0.5 as determined by titration. Two days post-transduction, cells were treated with 500 ng/mL puromycin and were selected for 5-7 days to enrich for infected cells.

After selection, cells were seeded at a density of 1.08×10^5^ cells/cm^2^ on 15 cm plates and left to incubate overnight. The next day, the media was replaced with either DMEM + 10% FBS (serum condition) or optiMEM (serum-starved condition) and cells were incubated overnight. Approximately 4-6 hours prior to sorting, cells were treated with 2.5 μg/mL BODIPY™ FL LDL (Thermo Fisher) in optiMEM. Cells were trypsinized and sorted into four bins (bottom 12.5%, bottom 12.5-37.5%, top 37.5-12.5%, and top 12.5%) or two bins as specified in the text using a BD FACSAria flow cytometer. After sorting, genomic DNA was isolated from cells using the Purelink Genomic DNA mini kit (Thermo Fisher), and up to 20 μg of genomic DNA per sample was used to amplify the U6-3’ to gRNA hairpin region with different in-line barcodes. PCR2 was performed to add full-length Illumina sequencing adapters using internally ordered primers with equivalent sequences to NEBNext Index Primer Sets 1 and 2 (New England Biolabs). All PCRs were performed using Q5UltraII polymerase (NEB). Pooled samples were sequenced using NextSeq (Illumina), using 75-nt reads and collecting greater than 100 reads per gRNA in the library.

The library prep primers were as follows:

PCR1:

U6_Bc_r1seq_halftail (24 distinct versions of this primer with staggered-length in-line barcodes denoted here as NNNNN were used)

5’ ACTCTTTCCCTACACGACGCTCTTCCGATCT NNNNN GGAAAGGACGAAACACCG 3’ gRNAFE_r2seq_halftail

5’ GACTGGAGTTCAGACGTGTGCTCTTCCGATCTGCCTTATTTAAACTTGCTATGCTGT 3’

PCR2:

r1seq_fulltail

5’ AATGATACGGCGACCACCGAGATCTACACTCTTTCCCTACACGACGCTCTTC 3’

r2seq_fulltail (up to 8 distinct indexed versions of this primer were used to maximize pooling)

5’ CAAGCAGAAGACGGCATACGAGATNNNNNNNNGTGACTGGAGTTCAGACGTGTGCT 3’

CRISPR-Cas9 screening analysis was performed using MAGeCK RRA v0.5.9.2 paired analysis comparing top and bottom sorted bins and normalized to non-targeting control gRNAs present in the library.

For the small molecule CRISPR-Cas9 screens, HepG2 Cas9NG-P2A-mChe cells containing the KO-Library 4 were seeded at a density of 1.08×10^5^ cells/cm^2^ on 10 cm plates in 3 replicates per condition and left to incubate overnight. The next day, the media was replaced with optiMEM (serum-starved condition) containing either control (DMSO) or one of eight inhibitors (see below) and cells were incubated overnight. Approximately 4-6 hours prior to sorting, cells were treated with 2.5 μg/mL BODIPY™ FL LDL (Thermo Fisher) in optiMEM. Cells were trypsinized and sorted into two bins (bottom 30% and top 30%) using a BD FACSAria flow cytometer. After sorting, genomic DNA was isolated from cells and PCR and sequencing was performed as described above. Small molecule CRISPR-Cas9 screening analysis was performed using MAGeCK RRA v0.5.9.2 paired analysis comparing top and bottom sorted bins and normalized to non-targeting control gRNAs present in the library. A default number of permutations were performed except when indicated otherwise in figure legends.

NGI-1 (Cayman) 5μM
ATN-224 (Cayman) 40 μM
KX2-391 (Cayman) 2.5 μM
Pyripyropene A (Cayman) 10 μM
Fatostatin (Cayman) 2.5 μM
MHY1485 (Cayman) 10 μM
GW3965 (Cayman) 0.02 μM
Rocaglamide A (MedChemExpress) 0.01 μM
Simvastatin (Cayman) 0.5 uM

#### Cloning and testing of individual gRNAs

Oligonucleotides including protospacer sequences (**Supplemental Table 21**) were ordered in the following format:

GGAAAGGACGAAACACCG [19-20-bp protospacer—remove initial G for any 20-bp protospacer with one natively] GTTTAAGAGCTATGCTGGAAAC were amplified by PCR to create homology arms and cloned into a lentiCRISPR-v2-FE-PuroR backbone through NEBuilder HiFi DNA assembly as described above. To make KO cell lines, the gRNA constructs were packaged into lentivirus and transduced into HepG2-Cas9NG-P2A-mChe cells seeded at 4×10^4^ cells/cm^2^ on 48-well plates in two replicates with 8 μg/mL of polybrene. Two days post-transduction, cells were treated with 500 ng/mL puromycin and were selected for approximately one week.

HepG2 Cas9NG-P2A-mChe KO cell lines were seeded 1:1 with HepG2 WT cells to achieve a total density of 1.08×10^5^ cells/cm^2^ on a 96 well plate in at least two technical replicates of two biological replicates per serum condition and incubated overnight. The next day, the media was replaced with either DMEM + 10% FBS (serum condition) or optiMEM (serum-starved condition) and cells were incubated overnight. Approximately 4-6 hours prior to FACS analysis, cells were treated with 2.5 μg/mL BODIPY™ FL LDL in optiMEM. Cells were trypsinized and analyzed for presence of mCherry and LDL uptake using a Beckman CytoFLEX flow cytometer. LDL uptake of each KO cell line was normalized to the LDL uptake of the WT cells within the same well. Differential LDL uptake between KO and control cells was further normalized using data from the control sgCTRL KO line.

ICE analysis was performed using Sanger sequencing reads and significance cut-off value for decomposition of P < 0.001.

#### Small molecule analysis

For the individual small molecule analysis, HepG2 Cas9NG-P2A-mChe cells were seeded at a density of 1.08×10^5^ cells/cm^2^ on a 96-well plate in 3-26 replicates per condition and left to incubate overnight. The next day, the media was replaced with optiMEM (serum-starved condition) containing either control (DMSO) or one of 24 inhibitors (**Supplemental Table 22**) and cells were incubated overnight. Approximately 4-6 hours prior to analysis, cells were treated with 2.5 μg/mL BODIPY™ FL LDL in optiMEM. Cells were trypsinized, resuspended in FACS suspension media, and analyzed by flow cytometric analysis using a Beckman CytoFLEX flow cytometer. Differential LDL-C uptake between small molecules and control treatment was further normalized using data from DMSO treated cells.

#### Lectin flow cytometry analysis

HepG2 individual gene KO cell lines were cultured in DMEM + 10% FBS with or without 5 μM NGI-1 for 24 hours. Cells were then trypsinized and 2.5*10^5 cells were resuspended in FACS suspension media (DMEM, no phenol red (Thermo Fisher 31053028) + 2% FBS + 2mM EDTA) containing streptavidin-Dy488 (2.5 ug/mL) and a biotinylated lectin (ECL (5 ug/mL) or WGA (0.625 ug/mL)). Staining reactions were incubated for 20 minutes, protected from light. Cells were washed twice and resuspended in FACS suspension media followed by flow cytometric analysis using a Beckman CytoFLEX flow cytometer. Lectin binding of each KO cell line was normalized using data from the control sgControl line.

#### RNA-seq

HepG2 Cas9NG-P2A-mChe KO cell lines were seeded at a density of 2.1×10^5^ cells/cm^2^ on a 24 well plate with two biological replicates per serum condition and incubated overnight. The next day, the media was replaced with either DMEM + 10% FBS (serum condition) or optiMEM (serum-starved condition) and cells were incubated overnight. RNA was harvested from cells using the Qiagen RNEasy Plus mini kit or the Zymo Quick-RNA 96 kit, prepared for RNA-seq using the Lexogen QuantSeq-Pool kit, and sequenced using Illumina Nextseq at >1*10^6 reads/sample.

RNA-seq reads were mapped using the Quantseq 3’ mRNA mapping pipeline as described by Lexogen to GRCh38.p12. Briefly, reads were first trimmed using bbduk from the bbmap suite (v38.79)^107^ trimming for low quality tails, poly-A read-through and adapter contamination using the recommended parameters. Then, reads were mapped using the STAR aligner (v2.5.2b)^108^ with the recommended modified-Encode settings. Finally, HT-seq (v0.9.1) count was used to obtain per-gene counts^109^.

Within each cell line, we conducted differential expression analysis using DESeq2 1.26.0 to identify significantly differentially expressed genes for each gene KO with respect to the control condition.

*OTX1* knockdown analysis was performed using published genome-wide Perturb-seq data^78^. Relative magnitude of change in expression (z-score) of 8,248 genes in K562 cells with *OTX1* knockdown was used as input in co-essential GSEA analysis using gseapy v.0.10.448 with 10,000 iterations and 60 parallel processes as negative control. The modules were analyzed according to their normalized enrichment scores (NES), and GO terms from PANTHER version 14 were linked as described below.

#### CRISPRa Outcome and Phenotype (COP) screen design and execution

An oligonucleotide library was designed pairing the proximal promoter of each of 200 target genes (194-nt upstream of the annotated TSS and 10-nt downstream) with 10 distinct gRNAs each, including 8 on-target gRNAs and 2 non-targeting gRNAs. On-target gRNAs were chosen as the 8 gRNAs with lowest Combined Rank from the CRISPick CRISPRa mode^42^ after filtering gRNAs that (i) are 1-nt offset from a lower-ranked gRNA on the same strand; (ii) contain an MfeI recognition site since MfeI was used in cloning this library. Off-target gRNAs were chosen from the non-targeting gRNAs in the Calabrese library^42^.

The COP oligonucleotide library (**Supplemental Table 3**) was ordered in the following format:

GGAAAGGACGAAACACCG [19-20-nt spacer] gtttaagagctaggccAACAATTGTCAACAGACCATGCC [205-nt promoter] GCTAGCTTGAAGGGGACG

The oligonucleotide library was amplified by PCR to create homology arms by using the primers; 010415_sgRNA_60bp_fw:

TAACTTGAAAGTATTTCGATTTCTTGGCTTTATATATCTTGTGGAAAGGACGAAACACCG, and 031621_CRISPRArepPCR1A_rv:

TGGTGAGTAGGAGGAAGAGGAAGCGCTTCATGGTGGCACGTCCGTCCCCTTCAAGCTAGC. In order to introduce 15 nt random barcodes, two separate fragments were also amplified by PCR wherea CD5F2_DPE_CTE_Puro-containing plasmid was used as the template by using the primers; 031621_CRISPRArep_PCR1B_fw: GCTAGCTTGAAGGGGACG and 031621_CRISPRArep_PCR1B_rv: CTAAGACAGGCGCAGCCTCCGAGGGATGTGTACATTTGGATGCAGGTCGAAAGGCNNNNNNNNNNNNNNNTCAACTGAAGCAGAAGAGGT, 031621_CRISPRArep_PCR1C_fw: GCCTTTCGACCTGCATCC and 031621_CRISPRArep_PCR1C_rv: CGGTGTTTCGTCCTTTCCAC, respectively. These three amplicons were then cloned into a plasmid backbone produced with NheI-HF and AgeI-HF double-digest of pLenti_CD5F2_MCPp65_Puro through NEBuilder HiFi, upscaled to 100ul including 0.05 pmol library, and 0.1 pmol of each PCR amplicon. After treating with Plasmid-Safe (Lucigen), the circularized assembly product was linearized with Mfel-HF digest and then was eluted using QIAquick PCR purification kit (Qiagen). The linearized assembly reaction was further amplified by PCR using the following primers; 072121_CRISPRArep_PCR2A_fwnew: tattacagggacagcagagatccagtttggttaattaaTTCCTTGTCAACAGACCATGCC and 050415_SAMsgRNA_60bp_rv atttaaacttgctaggccCTGCAGACATGGGTGATCCTCA TGTTggcctagctcttaaac and cloned into NheI+AgeI-digested pLenti_CD5F2_MCPp65_Puro backbone to produce a high throughput pooled library bearing proximal promoters of 200 LDL-related genes, paired gRNAs, as well as the CRISPRa SAM component MCP-p65-HSF1 in the same vector.

HepG2 cells used in the COP assay were first co-transfected with a non-autonomous Tol2 vector bearing dCas9-10xGcn4-mCherry-BlastR cassette (p2T CAG dCas9-10XGcn4-P2A-mCherry BlastR) and a Tol2 transposase-expressing vector using lipofectamine 3000 (Thermo Fisher). One day post-transfection, the cells were treated with 5ug/mL blasticidin and were selected for approximately 5 days. The cells were further sorted twice using a BD FACSAria flow cytometer for mCherry expression to produce a highly enriched HepG2^dCas910x-Gcn4-mChe^ cell population. This cell line was then transduced with a SunTag transcriptional activator lentiviral vector (pLenti_EF1a-scFv-SBNO1-KBP250-NFE2L1-GSTGGT-KRT40-P2A-BFP_HygroR) to produce a cell line stably expressing CRISPRa; HepG2^dCas9-10xGcn4-mChe-scFv-Sbno1-Nfe2l1-Krt40-BFP^. These cells were subsequently treated with the lentiviral COP library at a multiplicity of infection (MOI) of ~0.5. Two days post-transduction, cells were treated with 500 ng/mL puromycin and were selected for approximately one week.

To build a library connecting barcodes and gRNAs, genomic DNA was isolated from cells using the Purelink Genomic DNA mini kit (Thermo Fisher), and up to 20 μg of genomic DNA per sample was used to amplify promoter-barcode and barcode-gRNA pairs as dictionary library samples separately in PCR by using different in-line barcodes including primers shown below:

##### Paired Promoter-Barcode NGS sample

031621_CRISPRApro_r1seq_2-4N:

CTTTCCCTACACGACGCTCTTCCGATCT(N)_2-4_TCCTTGTCAACAGACCATGCC

031621_IntPri_r2seq_1-3N:

GGAGTTCAGACGTGTGCTCTTCCGATCT(N)_1-3_TTTGGATGCAGGTCGAAAGGC

##### Paired Barcode-gRNA NGS sample

031621_CRISPRa_CD5F2_r1seq_2-4N:

CTTTCCCTACACGACGCTCTTCCGATCT(N)_2-4_CACCTCTTCTGCTTCAGTTGA

031621_CRISPRA_MS2hp_r2seq_1-3N: GGAGTTCAGACGTGTGCTCTTCCGATCT(N)_1-3_CATGTTGGCCTAGCTCTTAAAC

PCR2 was performed to add full-length Illumina sequencing adapters using internally ordered primers with equivalent sequences to NEBNext Index Primer Sets 1 and 2 (New England Biolabs). All PCRs were performed using Q5UltraII polymerase (NEB). Pooled samples were sequenced using NextSeq (Illumina), using 40-45-nt reads and collecting greater than 100 reads per construct in the library.

In parallel, total RNA (RNEasy Maxi kit, Qiagen) and genomic DNA (Purelink Genomic DNA mini kit) were isolated from each biological replicate. cDNA samples were synthesized from total RNA samples with reverse transcription by using sequence specific primer, 031621_CTE_RTprimer: CCTCCGAGGGATGTGTACA, targeting downstream of the 15nt barcode site. In order to quantify the transcript abundancy of barcodes which would eventually demonstrate the activity of the promoter library in response to their paired gRNAs, 10ug gDNA and 20ug cDNA were PCR amplified by using 8 different r1, and a constant r2 primer for each samples: 122320_CD5F2_3pr2seq: GGAGTTCAGACGTGTGCTCTTCCGATCTNNNGCCAATGCCATAATCCACCTCT 121016_IntPri_r1seqtail_Bc: CTCTTTCCCTACACGACGCTCTTCCGATCT(N)_2-6_TTTGGATGCAGGTCGAAAGGC

PCR2 was performed to add full-length Illumina sequencing adapters using internally ordered primers with equivalent sequences to NEBNext Index Primer Sets 1 and 2 (New England Biolabs). All PCRs were performed using Q5UltraII polymerase (NEB). Pooled samples were sequenced using NextSeq (Illumina), using 75-nt reads and collecting greater than 100 reads per gRNA in the library.

For phenotypic LDL-C uptake screening, COP library-containing HepG2^dCas9-10xGcn4-mChe-scFv-Sbno1-Nfe2l1-Krt40-BFP^ cells were seeded at a density of 1.08×10^5^ cells/cm^2^ on 10 cm plates and left to incubate overnight. The next day, the media was replaced with optiMEM (serum-starved condition) and cells were incubated overnight. Approximately 4-6 hours prior to FACS sorting, cells were treated with 2.5 μg/mL BODIPY™ FL LDL in optiMEM. Cells were trypsinized and sorted into 2 bins (bottom 30%, top 30%) using a BD FACSAria flow cytometer. To quantify the gRNA abundance in sorted cell populations, genomic DNA was isolated from cells using the Purelink Genomic DNA mini kit (Thermo Fisher), and up to 20 μg of genomic DNA per sample was used to amplify U6-3’ to gRNA-FE hairpin region with in-line barcodes including primer pairs, all of which are different for each sample, shown below:

101317_U6PE1_BcXr1: ACTCTTTCCCTACACGACGCTCTTCCGATCT NNNNN GGAAAGGACGAAACACCG 031621_CRISPRA_MS2hp_r2seq_1-3N: GGAGTTCAGACGTGTGCTCTTCCGATC(N)_1-3_CATGTTGGCCTAGCTCTTAAAC

PCR2 was performed to add full-length index primers, shown below. All PCRs were performed using Q5UltraII polymerase (NEB). Pooled samples were sequenced using NextSeq (Illumina), using 50-nt reads and collecting greater than 100 reads per gRNA in the library.

031317_10xr2seq_24bp_N70X_fw: CAAGCAGAAGACGGCATACGAGATNNNNNNNNGTGACTGGAGTTCAGACGTGTGCT

110717_PE1_fulltail: AATGATACGGCGACCACCGAGATCTACACTCTTTCCCTACACGACGCTCTTC

#### COP data analysis

The Promoter-Barcode-gRNA dictionary was built upon two sets of pair-end sequencing datasets. In the first sequencing data, each R1 read contains a promoter sequence (>20-nt) from the 200 genes and the corresponding R2 read contains a 15-nt variable barcode sequence. A barcode-promoter pair was considered valid if the barcode predominantly (in >90% of reads) pairs with a single promoter sequence. In the second sequencing data, each R1 read contains the reverse complement of a barcode sequence, and the corresponding R2 read contains the reverse complement of a 19-20-nt gRNA sequence from the 2000 gRNAs targeting the promoters of the 200 genes. Similarly, a barcode-gRNA pair was considered valid if the barcode predominantly (in >90% of reads) pairs with a single gRNA sequence. The barcode-promoter pairs and the barcode-gRNA pairs were merged and saved to the promoter-barcode-gRNA dictionary if the gRNA and the promoter were from the same gene and the Hamming distance between the barcodes was less than 2.

For the cDNA and gDNA reporter sequencing data, the 15-nt variable barcodes were extracted from R1 reads, and matched to the existing barcodes in the promoter-barcode-gRNA dictionary if the Hamming distance was less than 2. For each gRNA,the total number of reads for every matched barcode in the cDNA and gDNA data was aggregated and normalized to reads per million (RPM). The efficiency of the gRNA was calculated as the log2 fold change (LFC) of the cDNA to gDNA barcode count. Within each gene, the gRNA efficiency was proportionally scaled to the 0-1 range, where the gRNA with the lowest LFC had an efficiency of 0, and the gRNA with the highest LFC had an efficiency of 1.

gRNA and promoter features that were used as predictive variables in the linear regression model for gRNA efficiency include CRISPick onTarget efficacy (continuous), CRISPick onTarget rank (ordinal), CRISPick offTarget rank (ordinal), Deep SpCas9 score (continuous), TATA motif presence (nominal), InR motif presence (nominal), gRNA orientation (nominal), and cutsite transcription start site (TSS) offsite bins (nominal). For each gRNA, the cutsite TSS offset was defined as the distance from the TSS to the cutsite. cutsite TSS offset was further grouped into 10nt bins for cutsite within 80-150nt range downstream of the TSS. To ensure model consistency, the 1600 gRNAs were bootstrapped into 100 subsamples, each containing 1600 gRNAs with replacement. For each of the subsamples, a linear regression model was built using the above mentioned features as predictive variables and the gRNA efficiency as the response variable.

The LDL-C uptake gDNA sequencing data is made up of 3 replicates after quality control curation for technical data quality, and each replicate consists of a “top30” sample and a “bottom30” sample. Sequences were extracted from R1 reads and matched to the 2000 gRNA library. For each gRNA in the library, we recorded the matched read count for the “top30” sample and the “bottom30” sample, and calculated the “Top30/Bottom30” read count ratio. Pearson correlations were calculated for each replication pair on the “Top30/Bottom30” ratio. Read counts were selected as the input to the MAGeCK “test” module to calculate the log2 fold change (LFC) and the associated p-value of LDL-C uptake for each gene. To run the MAGeCK “test” module, we specified “Top30” samples (3 replicates) as treatment samples, “Bottom30” samples (3 replicates) as control samples, and we used control gRNAs as the normalization method.

#### Cloning and testing of individual COP gRNAs

Oligonucleotides including protospacer sequences (**Supplemental Table 21**) were ordered in the following format:

GGAAAGGACGAAACACCG [19-20-bp protospacer—remove initial G for any 20-bp protospacer with one natively] GTTTAAGAGCTAGGCCAACATG

Fifteen gRNAs were amplified by PCR to create homology arms and individually cloned into a pLenti-U6-2xMS2gRNA-MCPp65-PuroR backbone through NEBuilder HiFi DNA assembly as described above. To make CRISPRa cell lines, the gRNA constructs were packaged into lentivirus and transduced into HepG2 cells expressing dCas9-10xGcn4-mChe and scFv-Sbno1-Nfe2l1-Krt40-BFP, seeded at 4×10^4^ cells/cm^2^ on 6-well plates with 8 μg/mL of polybrene in at least two replicates per gRNA. Two days post-transduction, cells were treated with 500 ng/mL puromycin and were selected for approximately one week.

For the cell-surface protein staining, CRISPRa cells were washed, trypsinized, and 6.4*10^4 cells were resuspended in FACS suspension media containing one of four APC-antibodies:

APC anti-human CD36L1 (Biolegend)
APC anti-human CD132 (Biolegend)
APC anti-human CD71 (Biolegend)
APC anti-human CD222 (Biolegend)

Staining reactions were incubated on ice for 20 minutes and protected from light. Cells were washed twice and resuspended in FACS suspension media followed by flow cytometric analysis using a Beckman CytoFLEX flow cytometer to determine magnitude of activation of each cell surface protein. CRISPRa potency of each cell line was normalized to control wells that received a different gRNA and the same antibody.

#### Co-essentiality modules and enrichment

Modules were generated using DepMap 21Q2 Project Achilles data^51^. This data was altered to account for gene proximity effects using the method described by FIREWORKS^55^. The proximity corrected data was then converted to modules using an existing method^56^. Module enrichments for gene subsets were calculated from the hypergeometric distribution implemented in scipy v1.6.3^110^. Gene Ontology^111^ (GO) terms were generated from PANTHER version 14 along with the tool’s implementations of term enrichment and FDR control^112^. All GSEA analysis was performed with gseapy v.0.10.4^48^ with 10,000-100,000 iterations as negative control. Networks were constructed for all modules with a hypergeometric test FDR less than 0.01. t-Distributed Stochastic Neighbor Embedding (t-SNE)^113^ algorithm was used to reduce the dimensionality of the Achillies proximity corrected data, and the first and second principal components were used as the X and Y coordinates of the genes, respectively.

#### Exome burden analysis

UKB variant extraction is performed using Hail v0.2.66^114^. Variant allele frequencies were estimated from the Genome Aggregation Database (gnomAD v2.1)^115^. Variants were included with population maximum allele frequencies of <= 0.001 (Ensembl gnomAD plugin)^116^. Functional consequences and computational score predictions come from Variant Effect Predictor v99^117^ leveraging the dbNSFP v4.0 plugin^118^. Variant functional consequences terms were called deleterious if the VEP consequence indicated frameshift, nonsense, start lost, splice donor/acceptor disruption, or stop lost.

Individuals carrying multiple coding variants in the same gene were assigned the variant with the more impactful functional consequence (Deleterious, Missense). If multiple variants have the same worst functional consequence, the variant with the lowest population maximum allele frequency is selected. If these frequencies are the same, the variant with the greatest CADD^73^ score is chosen. Comparisons between carriers and non-carriers are performed with the Mann-Whitney U test utilizing scipy v1.6.3.^110^

Phenotypic adjustments are performed through ten-fold cross validation on a linear regression, subtracting feature weight associated with each covariate times the individual’s covariate from the original reported phenotype. Linear regressions were performed using scikit-learn v0.24.0.^119^

Clinvar analysis was restricted to all Pathogenic and Benign coding missense variants as reported in the May 2022 release.

Cox proportional hazard analyses were performed using lifelines v0.27.4^120^. [https://doi.org/10.21105/joss.01317]. Regressions were performed with presence of a rare missense variant and presence of an LOF variant as binary features. Most severe variant was determined using the same strategy previously mentioned. Exon coordinates were determined for genes of interest using MANE transcripts, with an additional 5nt retained up- and downstream of each coding region to capture splice acceptor and donor region variants. Gene-level VCF files were extracted from the UK Biobank WES joint-called pVCFs using bcftools. The VCF files were then normalized to flatten multiallelic sites and align variants to the GRCh38 reference genome. Variants located in NIST Genome in a Bottle “difficult regions” were removed from analysis, as were variants with a minor allele frequency greater than 0.1% in the UK Biobank cohort. Further filtering removed variants where more than 10% of samples were missing genotype calls^121^ and variants that did not appear in the UK Biobank cohort. To mitigate differences in sequencing coverage between individuals who were sampled at different phases of the UK Biobank project, variants were only retained in the final set if at least 90% of their called genotypes had a read depth of at least 10. All filtering took place by running bcftools through the Swiss Army Knife app on the UK Biobank Research Analysis Platform.

#### Significance Testing

Significance is reported based on a false discovery rate of 0.1 unless noted otherwise. False discovery rate corrections are done using the Benjamini-Hochberg procedure^122^ correcting for the number of independent tests performed. For comparisons involving multiple dependent tests (e.g. genes at different groupings of variants) we apply Simes Method^123^ on the group of hypotheses to represent a single independent test.

#### Gene overexpression and RAB10 pathway imaging analysis

To construct HepG2 cell lines with inducible gene overexpression, we first made a stable HepG2 rtTA expressing cell line by performing lentiviral infection with pLenti_rtTA-P2A-BFP_HygroR and sorting stably expressing cells for BFP expression. We then replaced the insert in the Tet promoter-containing vector pCW-Cas9-Blast (Addgene 83481) for LDLR-mCherry, RAB10-T2A-RABIF-P2A-mCherry, OTX2-P2A-mCherry, CSK-P2A-mCherry, or P2A-mCherry as a control. We made stable lines using lentiviral transduction into HepG2-rtTA-BFP cells using Blasticidin selection to select for cells that received the insert.

For LDL-C uptake experiments, we used the protocol above for individual gRNA testing, comparing mCherry+ gene-overexpressing cells with mCherry-wild-type control cells.

To examine the role of the RAB10 pathway in LDLR expression and LDL-C uptake, we infected HepG2 cells and HepG2-rtTA-BFP + Tet-LDLR-mCherry cells with sgRAB10, sgDENND4C, and sgCTRL gRNA lentivirus and selected for infected cells with Puromycin. To measure surface LDLR expression, we stained HepG2 KO cells with mouse anti-LDLR Clone C7 monoclonal antibody (BD 565641) and donkey anti-mouse DyLight649 (Jackson ImmunoResearch) for 20 min on ice followed by flow cytometric analysis using BD CytoFLEX. To track LDLR localization over time, we induced LDLR-mCherry expression in HepG2-rtTA-BFP + Tet-LDLR-mCherry KO cells using 2 μg/mL Doxycycline for the specified time followed by fluorescent imaging using a Zeiss Apotome microscope at 40x resolution. Images were processed using Zeiss ZEN software and ImageJ. We also performed flow cytometry on HepG2-rtTA-BFP + Tet-LDLR-mCherry KO cells induced to express LDLR-mCherry with 2 μg/mL Doxycycline for the specified time and chased with standard media for the indicated times.

#### RAB10 3’ UTR evaluation

We amplified the entire RAB10 3’ UTR sequence from HepG2 genomic DNA and inserted it immediately after the stop codon of GFP-RAB10 lenti (Addgene 130883). We also cloned an identical plasmid containing the minor allele of rs142787485. We performed lentiviral transduction of HepG2 with both plasmids, selecting to completion with Puromycin and performing flow cytometry using a BD CytoFLEX to measure GFP levels.

#### Mouse AAV8 shRNA experiments

All animal care and experimental procedures used in this study were approved by the Institutional Animal Care and Use Committee at National University of Singapore. Mice were housed under a 12-hour light-dark cycle with free access to water and normal chow diet. Male C57BL/6 littermates were obtained from *InVivos* at 6 weeks of age. For studies with special diet, 12-week-old mice were placed on western diet (TD.10885, Envigo) for a total period of 2 weeks.

To knock down hepatic *Rabif, Otx2 or Csk in vivo*, AAV8-scramble-shRNA (Vector Biolabs), AAV8-*Rabif*-shRNA (Vector Biolabs), AAV8-*Otx2*-shRNA (Vector Biolabs), or AAV8-*Csk*-shRNA (Vector Biolabs) were administered to eight-week-old C57BL/6 male mice via retro-orbital injection, with a dose of 6 x 10^11^ gc per mouse. Heparinized blood samples were taken at 2, 4 and 6 weeks post-injection for plasma lipid analysis, western diet-feeding started at 4 weeks post-injection. Mice were sacrificed at 6 weeks post-injection and livers and plasma were collected for further analysis.

Sequences of shRNAs used for *in vivo* gene targeting are as follows.

*Rabif*: GCAGACTTGTTGGTACTAATA
*Otx2*: GCTGTTACCAGCCATCTCAAT
*Csk*: GCCTTGAGAGAGAAGAAATTT

Blood was collected from tail tips after overnight fasting, and plasma was further isolated via centrifugation. Triglyceride and total cholesterol levels were measured using Infinity Triglycerides Reagent (Thermo Fisher) and Infinity Cholesterol Reagent (Thermo Fisher) respectively according to the manufacturer’s instructions. VLDL/LDL fraction of plasma was separated from HDL fraction using HDL and LDL/VLDL Quantitation Kit (Sigma) according to the manufacturer’s instructions.

Total RNA was extracted from mouse livers using Trizol reagent (Invitrogen). 1 μg of RNA was applied for reverse transcription with using Maxima H Minus cDNA synthesis system (Thermo). Quantitative real-time PCR (RT-PCR) was performed on a ViiA^TM^ 7 RT-PCR system (Applied Biosystems Inc.). Melting curve analysis was carried out to confirm the RT-PCR products. Expression data is presented after calculating the relative expression level compared to the housekeeping gene Rplp0.

The protocol for hepatic lipid extraction was adapted from previously published work^124^ with slight modifications. Briefly, to measure the lipid (Triglyceride and cholesterol) levels of liver, snap-frozen liver samples were weighed and homogenized in ten volumes of ice-cold PBS. Two-hundred microliters of the homogenate was transferred into 1,200 ul of chloroform:methanol (2:1; v/v) mixture followed by vigorous vortex for 30 seconds. One-hundred microliters of ice-cold PBS was then added into the mixture and mixed vigorously for 15 seconds. The mixture was then centrifuged at 4,200 rpm for 10 minutes at 4 degree. Two-hundred microliters of the organic phase (bottom layer) was transferred into a new tube and evaporated for dryness. Two-hundred microliters of 1% Triton X-100 in ethanol was used to dissolve the dried lipid with constant rotation for 2 hours. Triglyceride and cholesterol content were determined using the Infinity Triglycerides reagent (Thermo Scientific) and Infinity Cholesterol reagent (Thermo Scientific) respectively, based on manufacturer’s instructions.

#### Quantification and Statistical Analysis

Number of replicates can be found in the Figure legends or in the Methods Details of STAR methods. All figures show mean with standard error bars unless specified otherwise. Two-tailed t-tests and Mann-Whitney U-tests were performed to compare treatment and control groups as indicated in Figure legends. P-values are represented by asterisks (*p<0.05, **p≤0.01, ***p≤0.001). Statistical analysis and visualization were carried out in Graph Pad Prism version 9.3.1 and R version 3.6.3. Flow cytometric analysis was performed using FCS Express version 7.12 and FloJo version 10.6.2.

## Acknowledgments

The authors thank Grigoriy Losyev, Benjamin Holmes, Mandana Arbab, Allison James, and Benjamin Angulo for technical assistance. The authors acknowledge funding from 1R01HG008754 (D.K.G., R.I.S.), 1R21HG010391 (R.I.S., C.A.C.), R01HG010372 (J.D.F. and C.A.C.), American Cancer Society (R.I.S.), American Heart Association (R.I.S.), National Organization for Rare Diseases (R.I.S.), Qatar Biomedical Research Institute (R.I.S.), and a National University of Singapore Start-up grant (H.J.Y.). Figures created with BioRender.

## Author contributions

Conceptualization, Methodology, Writing – Original Draft and Writing – Reviewing and Editing: R.I.S., C.A.C., H.Y., M.C.H., J.D.F, E. A., T.Y., D.K.G. and G.L.; Investigation and Validation: R.I.S., H.Y., M.C.H, E. A., B.K., M.C. and S.B.; Software, Formal Analysis and Visualization: R.I.S., C.A.C., H.Y., M.C.H, J.D.F, E.A., T.Y., B.K., G.H.T.Y. and G.L.; Funding Acquisition and Supervision: R.I.S., C.A.C., H.Y., D.K.G.

## Competing interests

The authors report no competing interests.

## Figures

**Figure S1:**
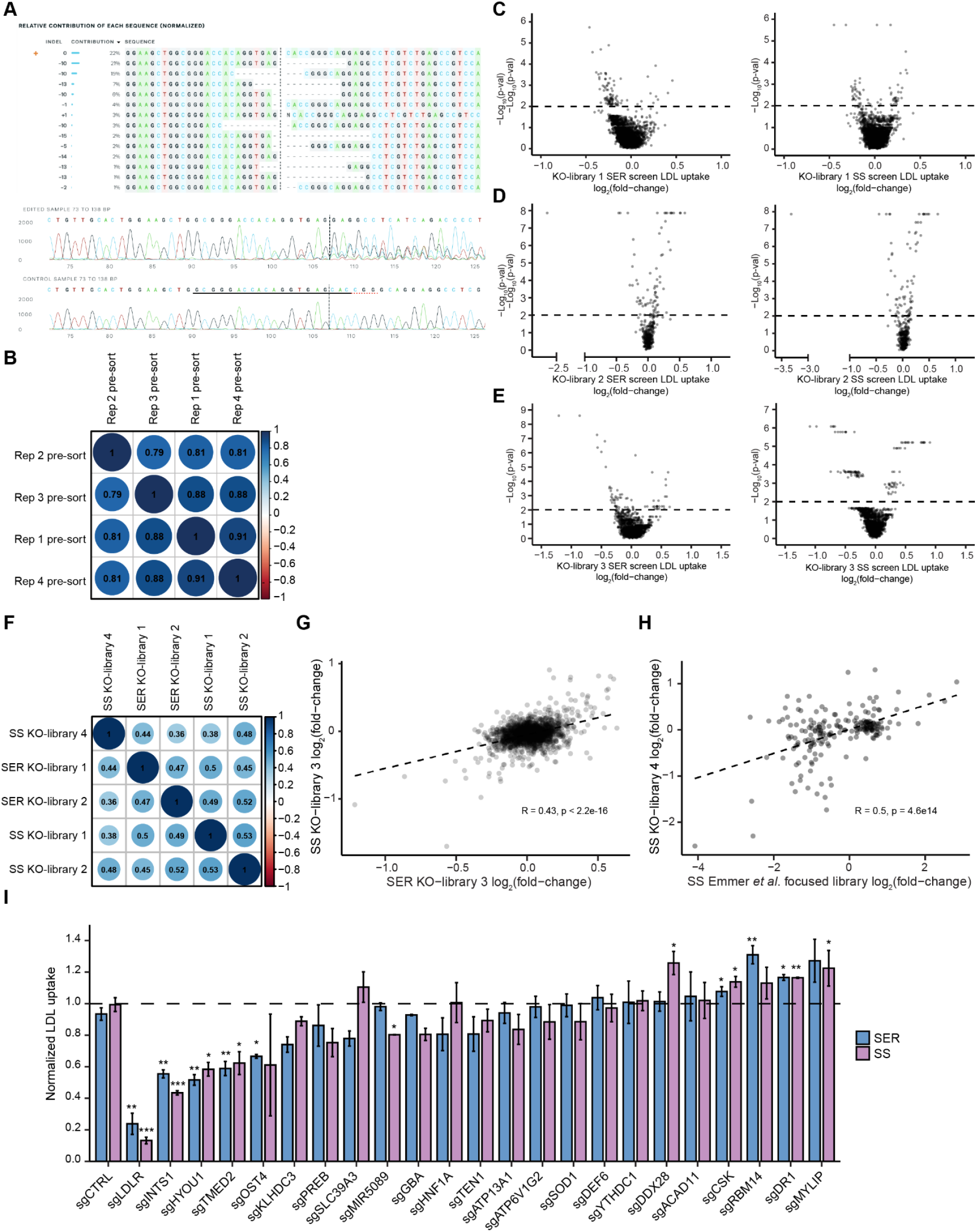
LDL-C uptake CRISPR-Cas9 knockout screening data, Related to Figure 1. A: Indel editing outcomes and trace files spanning the cut site from HepG2 cells with CRISPR-Cas9 knockout of LDLR and non-targeting control. Black underline = guide sequence, red dotted underline = PAM sequence, vertical dotted line = cut site. B: Spearman correlation analysis of the representation of all gRNAs in pre-sort cells in KO-Library 1 in SS condition. C-E: Volcano plots showing the MAGECK LDL-C uptake log2-fold-change (x-axis) and minimum MAGECK-Log10-p-value (y-axis) for the genes in KO-Library 1 (B), KO-Library 2, permutations = 100,000. (C), and KO-Library 3, permutations = 100,000. (D) in SER and SS conditions. F: Spearman correlation analysis of the log2-fold-change of the union set of 229 genes screened in KO-Library 1, KO-library 2, and KO-library 4 in SER and SS conditions. G: Spearman correlation analysis of the log2-fold-change of genes screened in the Emmer *et al*. focused library and KO-library 4 in SS condition. H: Spearman correlation analysis of the log2-fold-change of genes screened in KO-Library 3 in SER and SS conditions. I: Control-normalized LDL-C uptake of 23 individual gRNA knockout cell lines. n=2-4. Two-tailed t-test. P-values are represented by asterisks (*p<0.05, **p≤0.01, ***p≤0.001).

**Figure S2:**
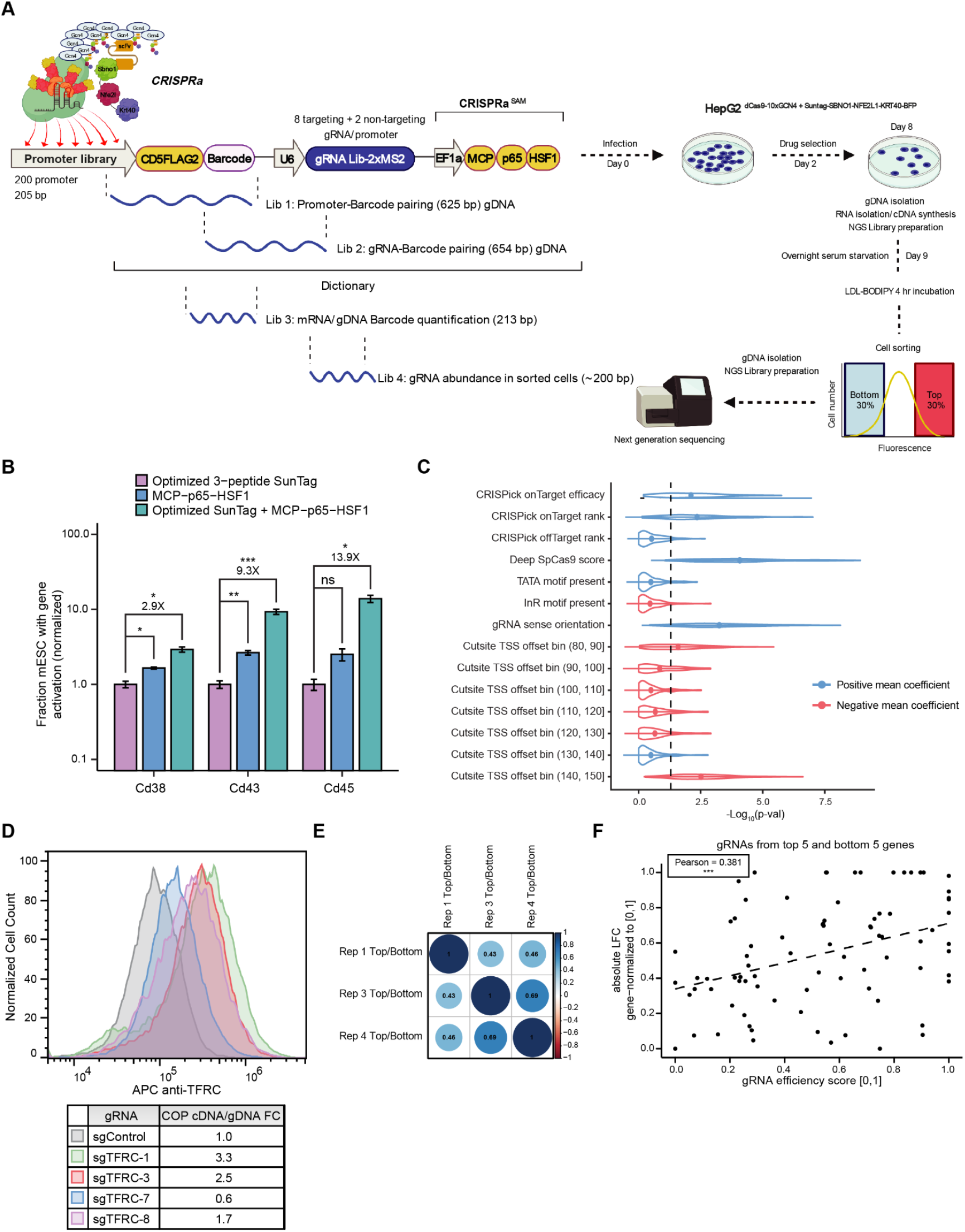
Analysis of CRISPRa Outcome and Phenotype screens, Related to Figure 2. A: Detailed schematic of CRISPRa Outcome and Phenotype (COP) screening workflow. See Figure 2A legend and text for details. B: Comparisons of the fraction of mESCs with activation (y-axis) of three target cell surface proteins (Cd38, Cd43, Cd45) in cells stably transfected with dCas9-10xGcn4, an MS2-gRNA targeting the target gene promoter, and ScFvGcn4-SBNO1-NFE2L1-KRT40 (purple), MCP-p65-HSF1 (blue), or both activation components (green). Cell fractions are normalized to results from ScFvGcn4-SBNO1-NFE2L1-KRT40 alone (purple). n=2-3. Two-tailed t-test. P-values are represented by asterisks (*p<0.05, **p≤0.01, ***p≤0.001). C: Distribution of −log10 p-values of COP gRNA-level reporter activity predictor terms in a linear regression model with bootstrapping. The mean and standard deviation of the p-values are shown with the superimposed point range graph from subsampling analysis. Predictor terms with positive mean coefficient across all subsamples are colored blue, and those with negative mean coefficient are colored red. The vertical dash line indicates the p=0.05 significance level. D: Flow cytometric analysis of cell surface antibody staining of TFRC in HepG2 cells with 4 individual CRISPRa gRNAs targeting the *TFRC* promoter or a control gRNA. Reporter cDNA:gDNA ratio is reported. E: Spearman correlation analysis of the ratio of representation of all CRISPRa library gRNAs in cells with top 30% vs. bottom 30% LDL-C uptake. F: Comparison of the CRISPRa reporter efficiency (x-axis) vs. phenotypic impact (y-axis) of gRNAs targeting the 10 genes with strongest LDL-C uptake phenotypes. Reporter efficiency is normalized gene-wise such that 0 indicates the lowest and 1 the highest cDNA:gDNA ratio of gRNAs targeting that gene. Phenotypic impact is normalized gene-wise such that 0 indicates the lowest and 1 the highest absolute LDL-C uptake log2-fold-change of gRNAs targeting that gene.

**Figure S3:**
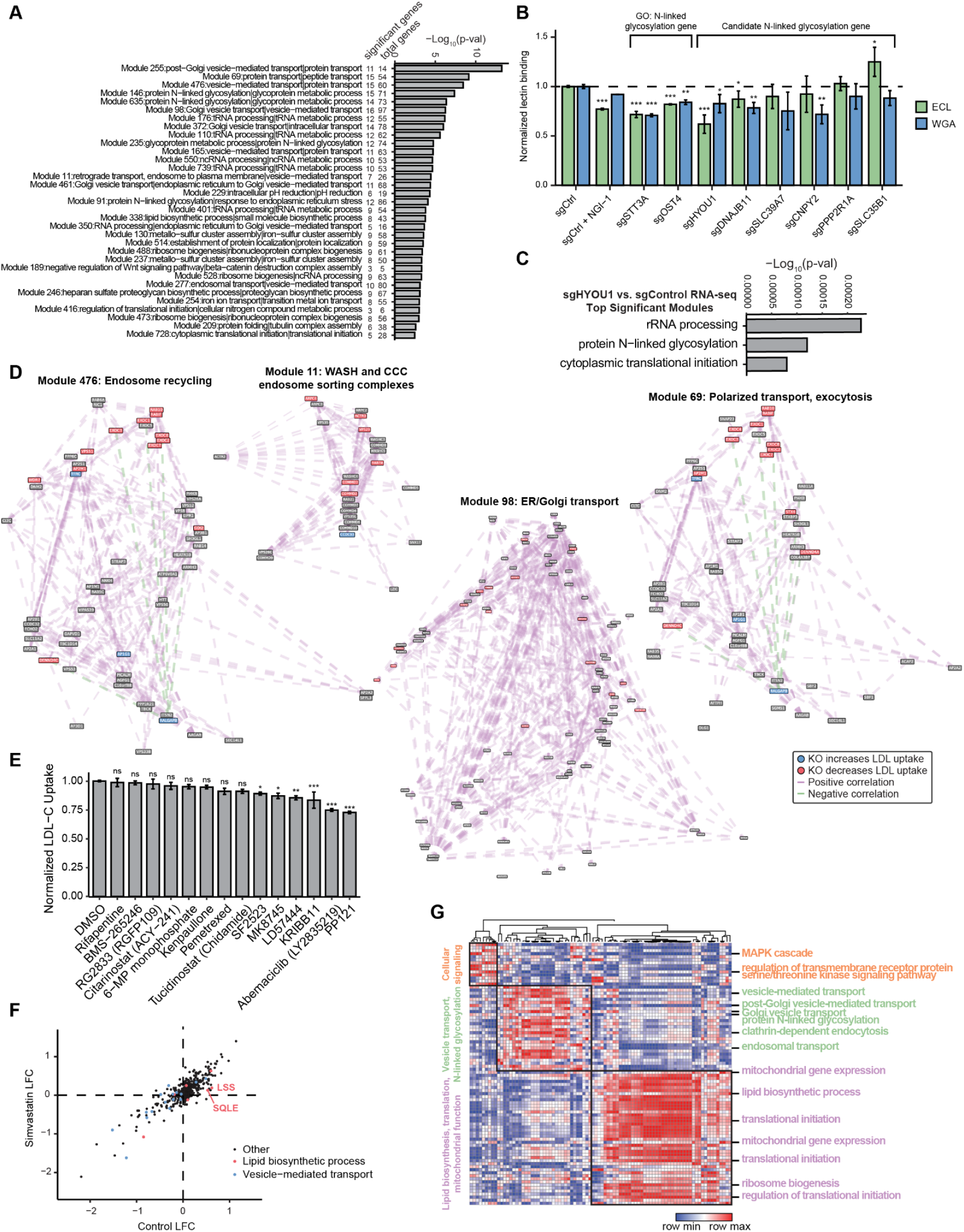
Co-essentiality analysis to identify LDL-C uptake-altering gene sets, Related to Figure 3. A: All co-essential modules with significant enrichment of CRISPR-KO genes (hypergeometric p-value) labeled with the module number and two most significantly associated GO terms for each module. B: Control-normalized flow cytometric binding of ECL (green) and WGA (blue) lectins in HepG2 treated with the indicated gRNAs and NGI-1. Two tailed t-test. n=3-5. C: Co-essential module enrichment among sgHYOU1 RNA-seq upregulated genes. D: tSNE plots based on PCA of modules 69 (polarized transport, exocytosis), 98 (ER/Golgi transport), 11 (WASH and CCC endosome sorting complexes), and 476 (endosome recycling). Genes that decrease LDL-uptake when knocked out are labeled red, and genes that increase LDL-uptake when knocked out are labeled blue. Genes with pairwise Pearson correlation of >0.2 (purple) or <-0.2 (green) are connected with dashed lines. E: Mean control-normalized LDL-C uptake fold-change of HepG2 cells treated with DMSO (control) or one of 14 small molecules. Dunnett’s multiple comparisons test. n=3-8. P-values are represented by asterisks (*p<0.05, **p≤0.01, ***p≤0.001). F: Comparison of LDL-C uptake log2-fold-change for 522 genes targeted by CRISPR-KO in the presence of DMSO (x-axis) and Simvastatin (y-axis). Genes in co-essential modules annotated to be involved in lipid biosynthetic process are shown in red and those involved in vesicle-mediated transport are shown in blue. G: Heat map displaying the Pearson correlation between every pair of co-essential modules from co-essential GSEA analysis of LDL-C uptake CRISPR-KO screening of 522 genes in the presence of eight small molecules. Modules are labeled with the most associated GO term and are organized by hierarchical clustering on the x-axis and K-means clustering (k=3) on the y-axis.

**Figure S4:**
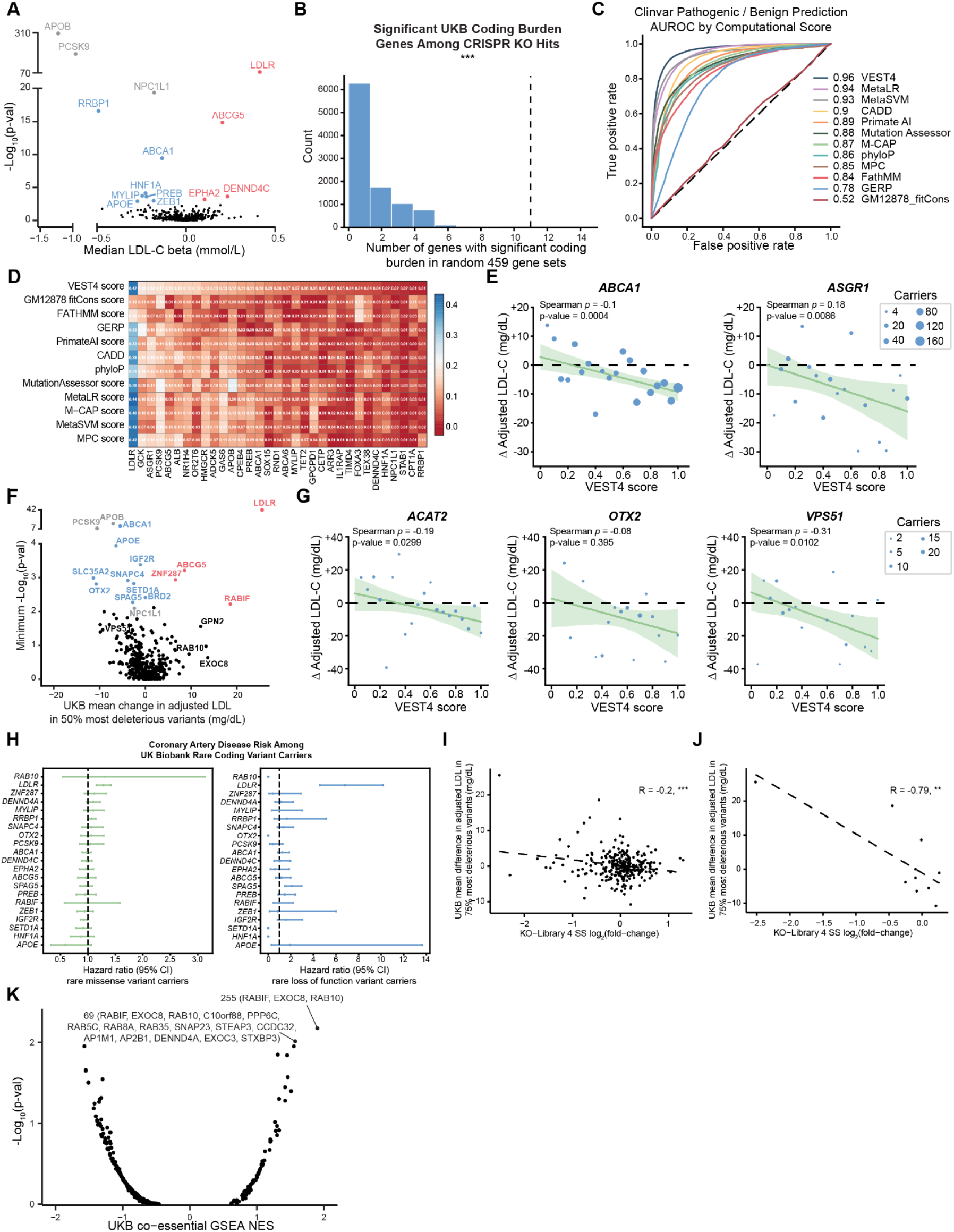
Gene- and co-essential module-level coding burden analysis results, Related to Figure 4. A: Volcano plot showing the median LDL-C beta (x-axis, mmol/L) and −Log10-p-value (y-axis) in the pLOF-restricted burden analysis for 459 genes with significant effects on LDL-C uptake in the CRISPR-KO screen as well as 3 additional well-known LDL-C burden genes (gray). Genes significantly associated with increased (red) and decreased (blue) LDL-C levels in either burden analysis are highlighted. B. Bar chart displaying the number of genes significantly associated with serum LDL-C upon 10,000 random selections of 459-gene sets from the pLOF-restricted burden analysis. 0/10,000 selections have at least as many significant genes (11) as when the 459 CRISPR-KO-significant genes are analyzed. C: AUROC comparison of 12 deleteriousness methods using ClinVar data. Computational scores are applied to all variants reported as Pathogenic or Benign as binary classification categories. D: Correlation comparison of 12 deleteriousness methods using UKB data for 33 genes with LDL-C burden from the pLOF-restricted approach. E: Correlation within UKB *ABCA1* and *ASGR1* variant carriers between VEST4 score and change in adjusted LDL-C as compared to non-carriers. VEST4 scores are binned to the nearest 5%, the bubble sizes represent the number of carriers within each bin, and a 95% confidence interval around the estimated linear regression line. F: Volcano plot showing the mean adjusted LDL-C difference (x-axis, mg/dL) and −Log10-p-value (y-axis) in the VEST4 burden analysis for 417 genes with significant effects on LDL-C uptake in the CRISPR-KO screen as well as 3 additional well-known LDL-C burden genes (gray). Genes significantly associated with increased (red) and decreased (blue) LDL-C levels in the VEST4 burden analysis are highlighted as well as selected non-significant genes (black). G: Correlation within UKB *ACAT2, OTX2*, and *VPS51* variant carriers between VEST4 score and change in adjusted LDL-C as compared to non-carriers. VEST4 scores are binned to the nearest 5%, the bubble sizes represent the number of carriers within each bin, and a 95% confidence interval around the estimated linear regression line. H: Pearson correlation between impact on cellular LDL-C uptake and the effect of the top 75% most deleterious variants on adjusted LDL-C levels in UKB carriers among all genes with significant effects on LDL-C uptake in the CRISPR-KO screens. I: Pearson correlation between impact on cellular LDL-C uptake and the effect of the top 75% most deleterious variants on adjusted LD-C levels in UKB carriers among all genes with significant effects on LDL-C uptake in the CRISPR-KO screens and significant effects in the coding burden analysis. P-values are represented by asterisks (*p<0.05, **p≤0.01, ***p≤0.001). J: Volcano plot showing the UKB LDL-C rank co-essential GSEA normalized enrichment score (NES) (x-axis) and −Log10-p-value (y-axis) for 291 co-essential modules with a ClusterOne threshold (d=0.2). The two most enriched modules are highlighted along with leading edge genes for those modules.

**Figure S5:**
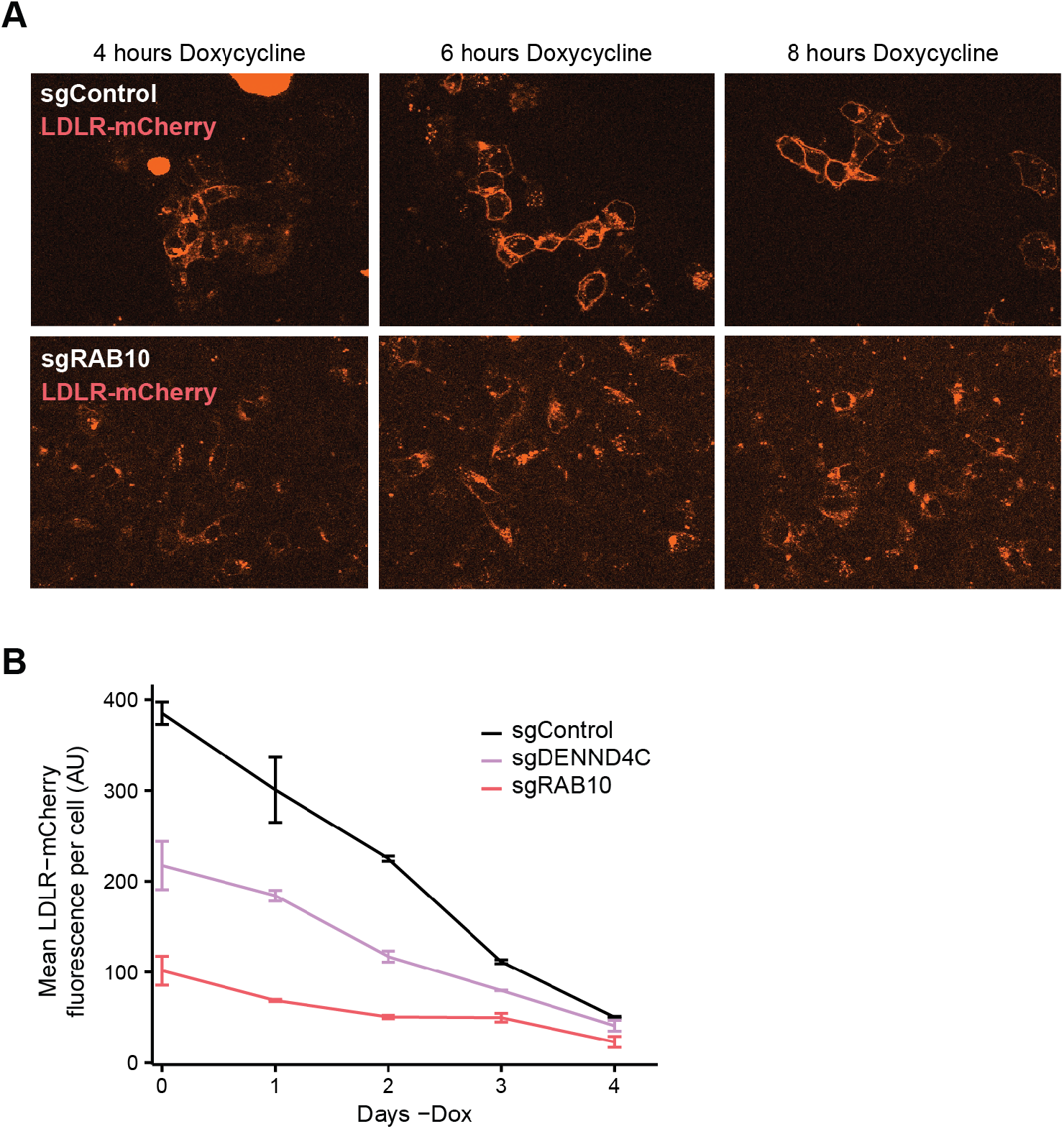
Timecourse analysis of LDLR-mCherry-expressing HepG2 in presence of RAB10 pathway knockouts, Related to Figure 5. A: Fluorescent microscopic images of HepG2 cells with inducible LDLR-mCherry expression and gene KO as indicated. LDLR-mCherry expression was induced 4, 6, or 8 hours prior to imaging as denoted. B: Flow cytometric expression of LDLR-mCherry after Dox removal in HepG2 cells with inducible LDLR-mCherry expression and gene KO as indicated.

**Figure S6:**
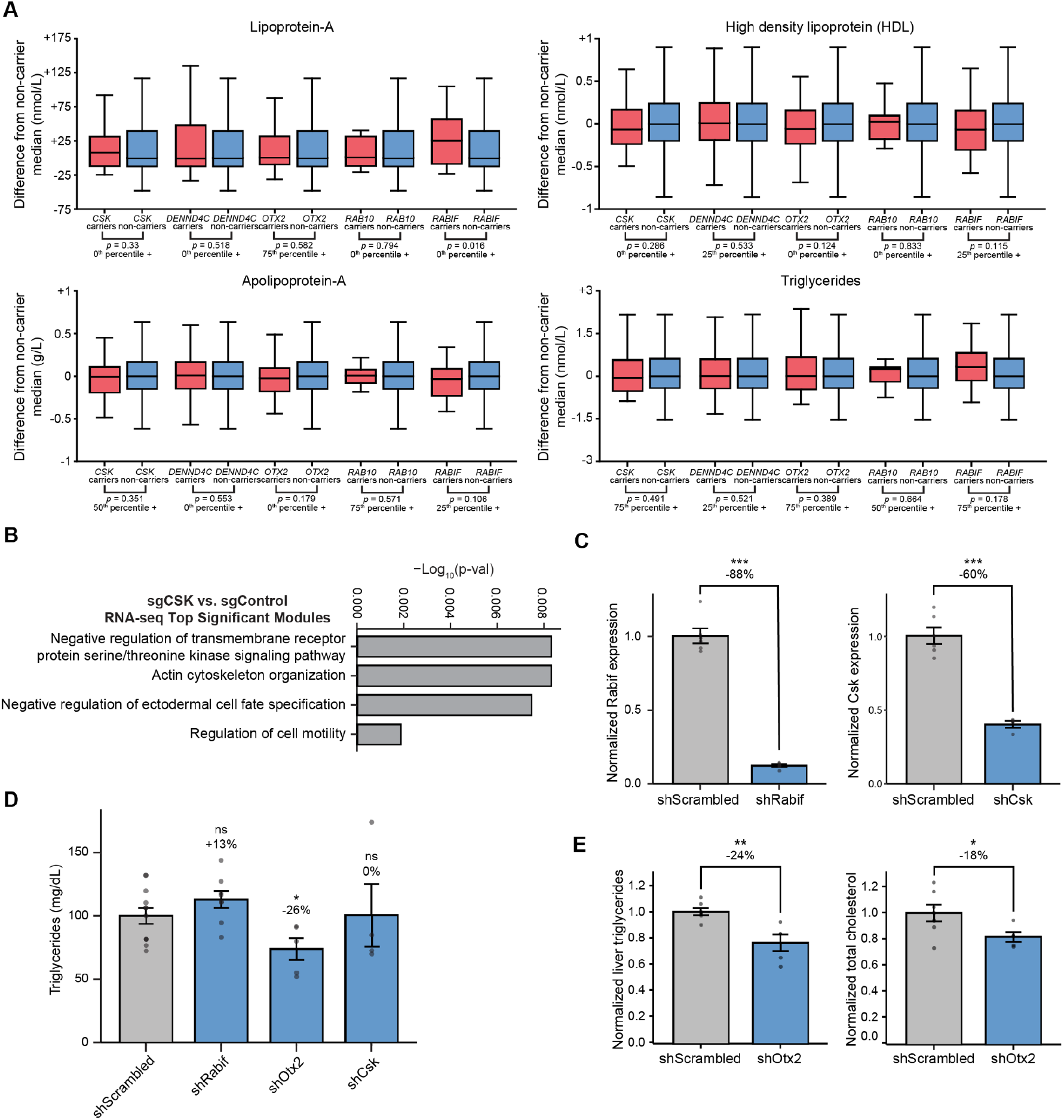
Evaluation of individual LDL-C uptake-altering genes by mouse AAV-mediated shRNA knockdown, Related to Figure 6. A: Analysis of Lipoprotein-A, HDL-C, Apolipoprotein-A, and Triglycerides in carriers (red) and non-carriers (blue) of coding variants in the top 50% of deleteriousness in *OTX2, CSK, DENND4C, RAB10*, and *RABIF*. P-values come from the Mann-Whitney U two-sided test using the minimum p-value from 0th+, 25th+, 50th+, or 75th+ percentile of deleterious variants noted below each gene’s comparison. B: Co-essential module enrichment among sgCSK RNA-seq upregulated genes. C: Whole mouse liver RT-qPCR analysis of target genes 6 weeks after AAV8 shRNA treatment. n=4-6. Two tailed t-test. P-values are represented by asterisks (*p<0.05, **p≤0.01, ***p≤0.001). D: Average triglyceride levels (mg/dL) in mice treated with the designated AAV8 shRNA 4 weeks post-treatment. Average difference compared to shScrambled and p-value are noted. n=4-12. Two tailed t-test. E: Whole mouse liver normalized triglycerides and normalized total cholesterol in mice treated with the designated AAV8 shRNA 6 weeks post-treatment. Two tailed t-test. n = 5-7.

**Figure S7:**
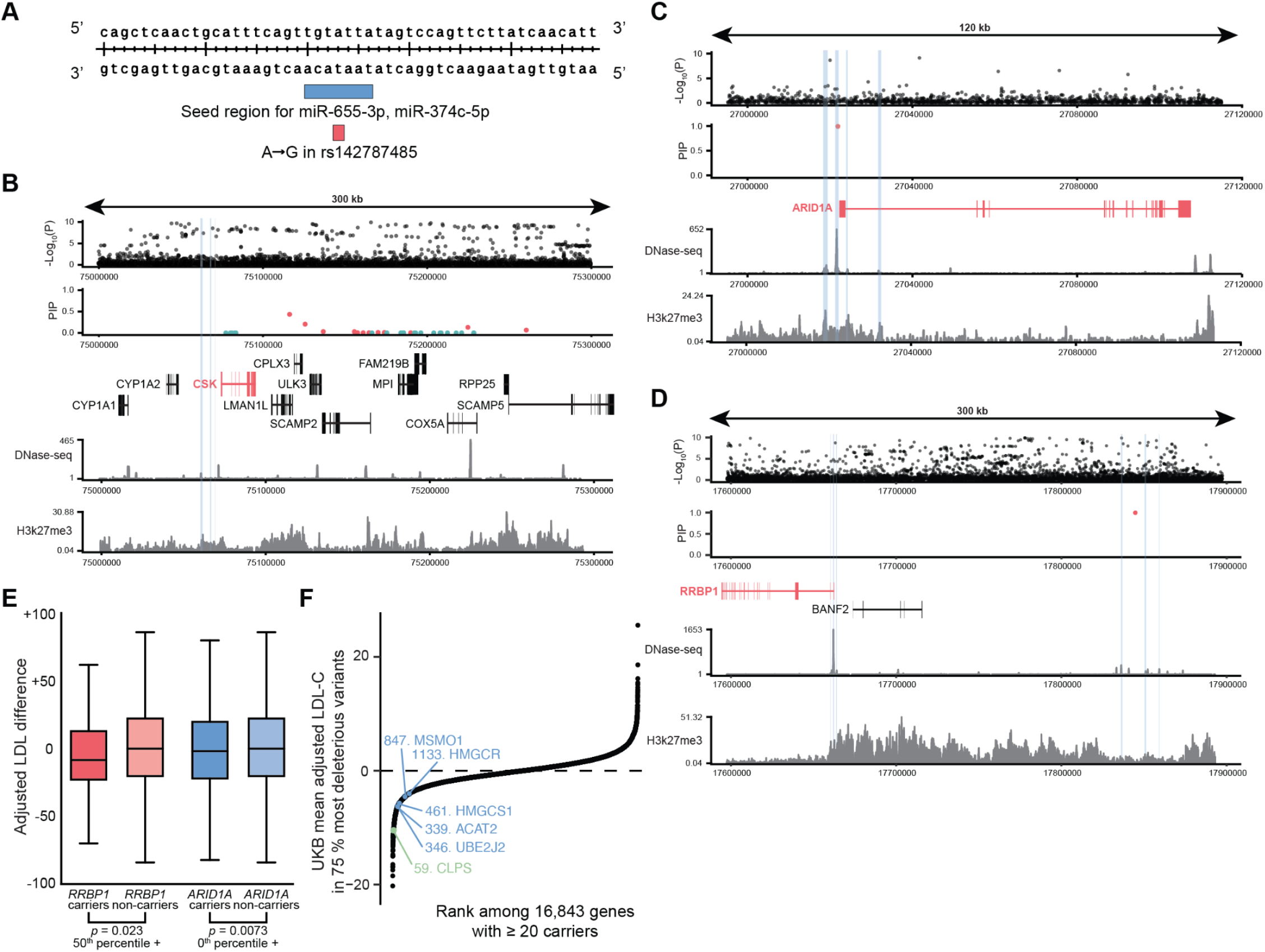
Examples of common variants surrounding genes with LDL-C burden, Related to Figure 7. A: Diagram of the rs142787485 (red) locus located within the RAB10 3’ UTR. The predicted binding site of two microRNA species is indicated in blue. B-D: Fine-mapping of the GWAS signal at the *CSK* (B), *ARID1A* (C), or *RRBP1* (D) locus (black: LDL-GWAS variants, red: 95% confidence set, teal: 99% confidence set). Signal tracks represent chromatin accessibility from DNase-seq or H3K27me3 histone modifications from ChIP-seq. Blue bars represent ABC predicted enhancer connections with *CSK* (B), *ARID1A* (C), or *RRBP1* (D). E: Box plots showing ARID1A and RRBP1 UKB LDL difference at cutoff of maximal significance F: Dot plot showing the rank in mean carrier adjusted LDL-C for the 75% most deleterious variants for 16,483 genes with 20 or more carriers. Select genes in the lipid biosynthesis co-essential module (module 338) are shown in blue, and CLPS is shown in green. Of these genes, only *HMGCR* shows a significant association with serum LDL-C in the genome-wide burden analysis.

## Supplemental Tables

**Supplemental Table 1**: RNA-seq reads per million and analyzed gene sets

**Supplemental Table 2**: MAGeCK analysis of CRISPR-KO screens using KO-Libraries 1-4 in SER and SS conditions

**supplemental table 3**: COP reporter cDNA and gDNA and LDL-C uptake sorted population gRNA counts

**Supplemental Table 4**: MAGeCK analysis of CRISPRa screens

**Supplemental Table 5**: Co-essentiality modules and significance testing from CRISPR-KO and UKB coding burden analyses

**Supplemental Table 6**: Small molecule CRISPR-KO screen MAGeCK LFC values

**Supplemental Table 7**: Co-essentiality GSEA results for small molecule CRISPR-KO screens

**Supplemental Table 8**: Co-essentiality GSEA clustering from small molecule CRISPR-KO screens

**Supplemental Table 9**: Combined significant genes from UKB collapsing coding burden analyses

**Supplemental Table 10**: pLOF-focused UKB collapsing coding burden analysis

**Supplemental Table 11**: VEST4 deleteriousness-thresholded UKB collapsing coding burden analysis

**Supplemental Table 12**: Cox proportional hazard regression results

**Supplemental Table 13**: Co-essential GSEA from VEST4 ranked burden genes, ClusterOne d=0.8

**Supplemental Table 14**: Co-essential GSEA from VEST4 ranked burden genes, ClusterOne d=0.2

**Supplemental Table 15:** sgOTX2 RNA-seq GSEA

**Supplemental Table 16**: K562 ChIP-seq and Perturb-seq

**Supplemental Table 17**: CRISPR-KO Library 1 gRNA information

**Supplemental Table 18**: CRISPR-KO Library 2 gRNA information

**Supplemental Table 19**: CRISPR-KO Library 3 gRNA information

**Supplemental Table 20**: CRISPR-KO Library 4 gRNA information

**Supplemental Table 21**: Individual gRNA oligos

**Supplemental Table 22:** Small molecule dosing and pathway targets

## Notes

https://github.com/j-fife/hamilton-et-al

### Competing Interest Statement

The authors have declared no competing interest.

